# The actin motor protein MYO10 facilitates post-entry spread of respiratory syncytial virus

**DOI:** 10.64898/2026.03.29.715182

**Authors:** Xiaoyu Zhang, Isra Nurhassen, Martina Friesland, Vanessa Milke, Felix Bender, Katharina Maser, Martin Wetzke, Gesine Hansen, Thomas Zillinger, Chris Lauber, Thomas Pietschmann

## Abstract

Human respiratory syncytial virus (RSV) is a leading cause of severe lower respiratory tract infections in infants. However, host factors that influence disease severity remain incompletely defined. While clinical risk factors are known, identifying genetic susceptibility has been challenging. In this study, we combined human genetics with functional virology to identify host factors that modulate RSV infection and spread.

Starting from a cohort of infants hospitalized with severe RSV disease, we prioritized rare coding variants present in homozygous form and predicted to cause strong functional impairment, and selected candidate genes for mechanistic follow-up. Functional interrogation of 23 candidates by CRISPR/Cas9 knockout screening in human lung epithelial cells identified unconventional myosin-X (MYO10), encoding the actin-based motor protein myosin-X, as a critical host factor for RSV. Genetic disruption or siRNA-mediated depletion of MYO10 significantly reduced RSV infectivity, with the strongest effects at post-entry stages of the viral life cycle.

Loss of MYO10 impaired filopodia formation, cell migration, and wound healing, leading to altered cell–cell connectivity and restricted viral dissemination. MYO10 depletion reduced both short-range cell-to-cell transmission and longer-distance extracellular spread, resulting in fewer infected cells and diminished accumulation of progeny virus in culture supernatants. In contrast, RSV entry, early gene expression, and interferon responses were unaffected.

Finally, a rare homozygous MYO10 motor-domain variant (rs7737765; H148Y), enriched in severe cases, also reduced RSV replication in cell culture—opposite to expectations for a risk allele—yet underscoring biological relevance and suggesting that MYO10 variation may influence disease in vivo through additional effects on epithelial function.

**Importance:** Respiratory syncytial virus (RSV) is a major cause of severe respiratory illness in infants, yet it remains unclear why some children develop more serious disease than others. In this study, we combined patient genetic data with laboratory experiments to identify host factors that influence how RSV spreads in the lung. We found that the human protein MYO10, which mediates the formation of small cell protrusions, plays a key role in enabling the virus to spread from cell to cell. When MYO10 was disrupted, viral spread was strongly reduced, even though early steps of infection were unaffected. Interestingly, a rare genetic variant in MYO10 found in patients also altered viral replication, highlighting potential clinical relevance. These findings provide new insight into how host cell architecture contributes to RSV infection and suggest that targeting host pathways involved in viral spread could complement existing antiviral strategies.

## Introduction

RSV is the leading viral cause of acute lower respiratory tract infections (ALRI) in infants and young children worldwide (Mazur, Caballero et al. 2024). Despite major advances in prevention, including long-acting monoclonal antibodies and maternal vaccination (Calvo and García-García 2025, Fly, Stultz et al. 2025, Madhi, Simões et al. 2025), RSV remains a substantial global disease burden, with millions of hospitalizations annually and significant mortality, particularly in low- and middle-income countries (Wang, Li et al. 2024). Severe RSV disease in infants is characterized by extensive epithelial damage, airway obstruction, and impaired gas exchange. However, the host determinants that predispose to severe disease remain incompletely understood.

Established clinical risk factors for severe pediatric RSV infection include prematurity, congenital heart disease, chronic lung disease, immunodeficiencies, Down syndrome and neuromuscular disease (Mazur, Caballero et al. 2024). However, the majority of infants hospitalized with RSV are born at term and lack identifiable comorbidities, indicating that additional host-specific factors contribute to disease severity. Age-dependent vulnerability, variations in airway structure and immune maturation likely play a role. Moreover, accumulating evidence suggests that host genetic variation may also influence RSV pathogenesis. Large genome-wide association studies (GWAS), however, have failed to identify common single nucleotide polymorphisms (SNPs) robustly associated with RSV hospitalization or disease severity (Egeskov-Cavling, van Wijhe et al. 2024, Johnson, Chelysheva et al. 2024). This suggests that genetic susceptibility to severe RSV disease may not be driven by common variants of large effect, but rather by rare or moderately penetrant variants that affect key host pathways and escape detection by conventional GWAS approaches.

Transcriptomic studies have provided complementary insights, revealing host gene expression signatures associated with severe RSV infection, including pathways linked to inflammation, epithelial integrity, and immune cell recruitment (Flerlage, Crawford et al. 2023, Zivanovic, Öner et al. 2023, Pastey and Lupfer 2025). While these studies highlight biological processes correlated with disease severity, they do not directly establish causal host factors that modulate RSV infection or spread. Moreover, genetic variants that alter protein function without necessarily changing transcriptional responses may critically shape host-virus interactions but remain underexplored.

Beyond viral entry and replication, the efficiency with which RSV spreads within the airway epithelium and is cleared from the respiratory tract is increasingly recognized as a major determinant of disease outcome. RSV dissemination occurs through a combination of direct cell-cell transmission and extracellular spread. These processes depend on dynamic remodeling of the actin cytoskeleton and membrane architecture (Mehedi, McCarty et al. 2016, Rentzsch, Witten et al. 2019, Kieser, Granoski et al. 2023, Zhang, Lin et al. 2024). Actin-driven cellular protrusions, including filopodia, can facilitate intercellular contacts, transport of viral material, and localized viral release, enabling rapid propagation across epithelial layers while partially shielding the virus from neutralizing antibodies (Zhang, Lin et al. 2024). In parallel, effective clearance of RSV from the airways relies on intact epithelial polarity, coordinated cell migration, and mucociliary function. Disruption of epithelial architecture or cytoskeletal dynamics can impair ciliary beating and mucociliary clearance, thereby prolonging viral persistence in the lung and exacerbating inflammation (Bustamante-Marin and Ostrowski 2017). Host factors that regulate cell shape, protrusion formation, migration, and epithelial organization may play a crucial role both RSV spread and clearance, yet these factors remain poorly defined.

Myosin-X, encoded by the MYO10 gene, is an unconventional actin-based motor protein with established roles in filopodia formation, cell migration, and membrane-cytoskeleton coupling (Liu and Cheney 2012, Courson and Cheney 2015). MYO10 localizes to filopodial tips and facilitates the transport of cargos along actin bundles, contributing to cell-cell interactions, signaling, and directed movement. Dysregulation of MYO10 has been implicated in altered cell motility and invasive behavior in cancer and developmental processes (Courson and Cheney 2015, Yu, Lai et al. 2015, Tokuo, Bhawan et al. 2018, Mas, Cristella et al. 2025). In the context of viral infections, filopodia and related actin-driven structures have been shown to promote viral assembly, budding, and intercellular spread, including for filoviruses, where myosin-X colocalizes with viral nucleocapsids during actin-dependent transport (Schudt, Kolesnikova et al. 2013). RSV infection itself induces profound actin cytoskeleton remodeling and filopodia formation in epithelial cells (Mehedi, McCarty et al. 2016, Kieser, Granoski et al. 2023). However, the contribution of specific actin-based motor proteins to RSV dissemination has not been directly addressed.

In this study, we leveraged patient-derived genetic data to identify host factors that modulate RSV infection and spread. Building on the “Infection with Respiratory Syncytial Virus in Infants” (IRIS) cohort of infants hospitalized with severe RSV disease (Wetzke, Funken et al. 2022), we applied a homozygosity-based filtering strategy to prioritize rare coding variants predicted to strongly impair protein function. This approach focuses on variants with high biological impact that may contribute to disease susceptibility. Functional validation of shortlisted candidate genes using CRISPR/Cas9-based knockout screening in human lung epithelial cells identified MYO10 as a critical host factor for RSV infection *in vitro*.

Through genetic disruption and siRNA-mediated depletion of MYO10, we demonstrate that MYO10 facilitates RSV infection predominantly at post-entry stages. Loss of MYO10 impairs filopodia formation, cell migration, and wound healing, leading to altered cell–cell connectivity and reduced extracellular viral dissemination. Importantly, MYO10 depletion does not affect RSV entry or interferon-mediated antiviral responses, indicating a pro-viral role independent of canonical innate immune signaling pathways. Finally, functional analysis of a rare MYO10 missense variant identified in patients with severe RSV disease supports the clinical relevance of MYO10-mediated cytoskeletal regulation in RSV pathogenesis.

Together, these findings link human genetic variation to a host cytoskeletal regulator that governs RSV spread in vitro and potentially clearance-related processes in the airway epithelium, providing new insights into host determinants of severe RSV disease.

## Results

### IRIS cohort and homozygosity analysis

Given the incomplete picture on genetic variants that impact disease severity of RSV infections, we developed a systematic filtering pipeline to identify host genetic factors potentially contributing to severe RSV infection (Figure 1A). Considering the proven relevance of the interferon (IFN) system and interferon-controlled antiviral mechanisms for the course of viral infections, we focused our analysis on host gene regulating IFNs or regulated by IFNs. Building on previously published data on IFN regulated genes (material and methods), we analyzed 5,141 IFN-related genes polymorphisms that may modulate RSV infection. To minimize confounding effects from heterozygous variants whose phenotypic impact may be masked or partial, we first extracted all homozygous single nucleotide variants (SNVs) observed across these 5,141-IFN-related genes from the IRIS cohort participants (Wetzke, Funken et al. 2022). This initial set, designated as “level 0” in the schematic, comprised over 41,000 SNVs identified in 101 patients with severe RSV infection and mapping to 2299 genes. Variants were categorized by mutation types, including untranslated region (UTR), nonsynonymous missense (NSM), nonsynonymous nonsense (NSN), and nonsynonymous frameshift (NSF), and they are annotated relative to the Non-Finnish European (NFE) subset of the Genome Aggregation Data Base gnomAD (https://gnomad.broadinstitute.org).

**Figure 1.**
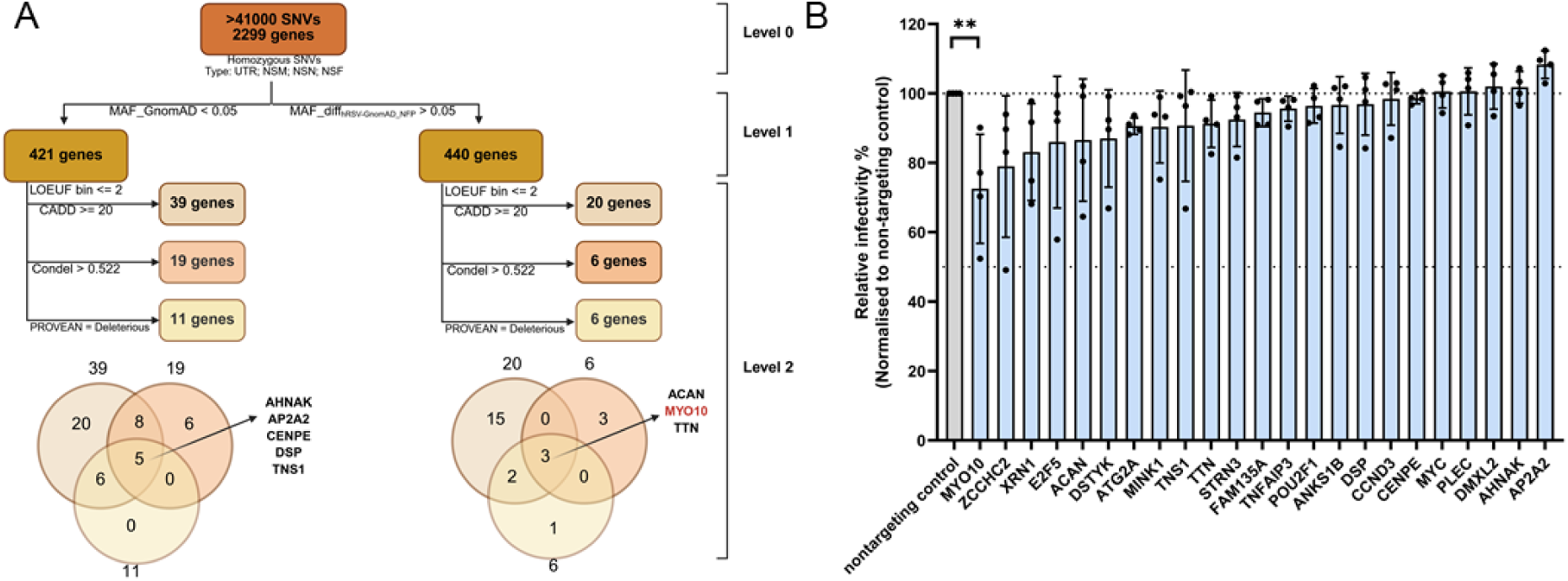
Homozygosity SNP filtering and CRISPR/Cas9 KO Screen. (A) Schematic overview of homozygosity analysis and single nucleotide polymorphism (SNP) filtering. CADD: Combined Annotation Dependent Depletion; Condel: CONsensus DELeteriousness Score; PROVEAN: Protein Variation Effect Analyzer; LOEUF: Loss-of-function observed/expected upper faction. (B) CRISPR Knock-Out (KO) screening of selected candidates. A549 cells stably expressing Cas9 were transduced with lentiviral particles packaging plasmids encoding individual sgRNAs with a dTomato reporter gene. Cells were infected 72 h post transduction with RSV-A-GFP (MOI 0.1) for 48 h and analyzed using flow cytometry. Data are presented as mean ± SD from four independent experiments. Statistical significance was determined by one-way ANOVA with multiple comparisons. ** p < 0.01; all other comparisons were not significant (p > 0.05) and are not indicated.

To further refine candidate variants, we implemented two parallel filtering strategies based on minor allele frequency (MAF). In the first, variants with MAF_gnomAD < 0.05 were retained. In the second, we selected variants with a MAF difference (MAF_diff) > 0.05 between the IRIS cohort and gnomAD. These complementary routes were chosen to account for both the low baseline hospitalization rate for severe RSV in the general population and the need to capture genetic differences enriched in the patient cohort. Each filtering branch reduced the candidate space to approximately 400 genes, collectively denoted as “level 1.”

Next, we applied multiple variant-effect prediction tools to assess functional impact, including CADD (Combined Annotation Dependent Depletion) (Rentzsch, Witten et al. 2019), Condel (CONsensus DELeteriousness score of non-synonymous single nucleotide variants) (González-Pérez and López-Bigas 2011), PROVEAN (Protein Variation Effect Analyzer) (Choi and Chan 2015) and LOEUF, a gene-level metric of intolerance to loss-of-function variants (Karczewski, Francioli et al. 2020). A Venn diagram was generated to illustrate the overlap between filtering subgroups, and genes harboring variants predicted to be deleterious by at least two tools were prioritized. This subset of candidates is designated as “level 2”.

Through this integrative filtering strategy, we ultimately identified 23 distinct genes containing high-confidence candidate SNVs for downstream functional validation. Given that Titin (TTN) was also found in the prioritization branch with low MAF, and thus occurred twice, in total homozygous variants in 23 distinct gene candidates met the inclusion criteria. Supplementary table S1 lists these candidate risk genes alongside the SNPs discovered by our analysis including the associated predicted variants-effects.

### CRISPR/Cas9 knockout screening of candidate genes

Following the identification of 23 candidate genes, we assessed their potential roles in RSV infection through a CRISPR/Cas9 knockout screen in A549 cells. Guide RNA sequences targeting each candidate gene were introduced into A549 cells stably expressing Cas9 by lentiviral gene transfer, yielding polyclonal knockout populations for each gene. The knockout cells were subsequently infected with RSV-A-GFP at an MOI of 0.1 for 48 hours, after which viral infectivity was quantified by flow cytometry. Because the guide RNA cassettes were delivered with a vector expressing a dTomato reporter, we first gated on dTomato-positive cells to control for lentiviral transduction efficiency, a proxy for overall knockout efficiency. RSV infection rate was quantified by the frequency of GFP-positive cells within the dTomato-positive cell population enabling direct comparison across candidate genes (Figure 1B, see supplementary Figure 1 for the gating scheme). From this primary screen, MYO10 emerged as the only gene whose knockout produced a statistically significant reduction in RSV infection. The minor, predicted deleterious allele in the MYO10 coding sequence (rs7737765; H148Y) was observed at a frequency of 16.8% in cases compared with 11.3% in gnomAD non-Finnish European controls, corresponding to an odds ratio of 1.59 (95% CI of 1.06-2.31) and indicating a modest enrichment (Fisher’s exact p = 0.0189) in the disease cohort (Supplementary table S2).

### MYO10 knockout and knock down limits RSV infection in A549 cells

To validate the effect observed in the primary CRISPR screen, we employed an orthogonal CRISPR-based strategy to generate MYO10-deficient A549 cells through CRISPR RNP delivery.

Sanger sequencing of single clones revealed a homozygous frame-shift deletion in clone 1H5 near the N-terminal region of MYO10, resulting in a premature stop codon (Figure 2A). Western blot analysis confirmed a markedly reduced level of MYO10 protein in clone 1H5 compared with A549wt cells; however, a low residual signal was still detectable (Figure 2B). This residual band may reflect nonspecific antibody binding or cross-reactivity, as the antibody used (Atlas Antibodies; rabbit polyclonal; Cat# HPA024223; RRID: AB_1854248) recognizes an epitope in the central region of MYO10 - between the IQ motifs and PH domains - which may share sequence similarity with other proteins of comparable molecular weight. Therefore, functional observations derived from clone 1H5 are interpreted as reflecting a substantial knockdown but potentially not a complete knockout.

**Figure 2.**
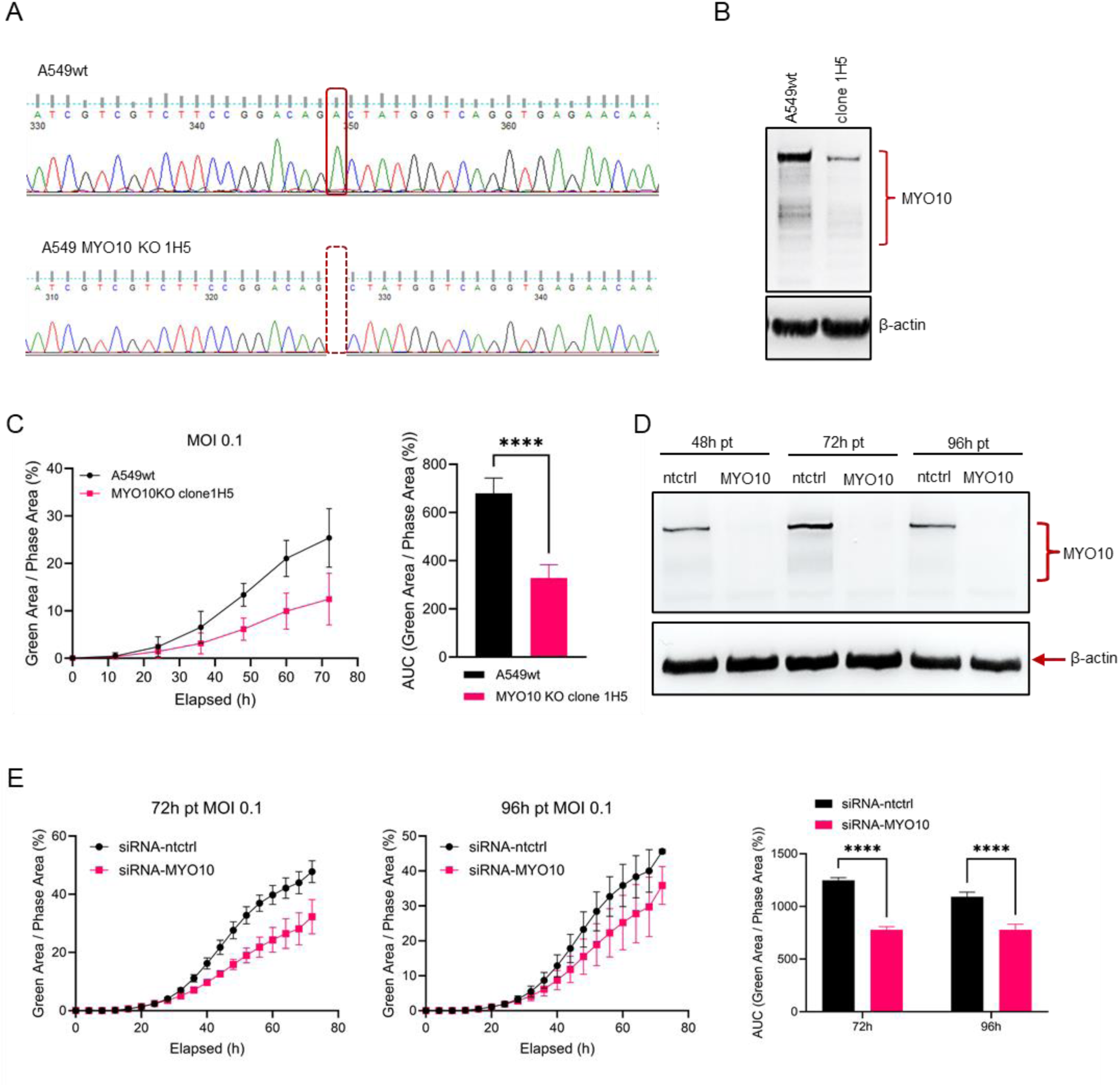
MYO10 knock out and knock down reduces RSV infectivity at late time post infection. (A) Sequencing alignment of A549 and MYO10 knock out single clone 1H5 created via CRISPR RNP transfection and single clone selection. Clone 1H5 possess one single nucleotide deletion, resulting in frameshift mutation. (B) MYO10 and β-actin expression in A549wt and MYO10 KO single clone 1H5 were measured by Western Blot. (C) MYO10 knock out single cell clone 1H5 together with control cell lines were infected with RSV-A-GFP at MOI 0.1 and monitored using Incucyte live-cell imaging (mean +/- SD, n=5). The area-under-curve (AUC) of green area per image normalized to phase area per image over 72h was calculated. Statistical significance was determined by one-way ANOVA with multiple comparisons (**** p < 0.0001). (D) MYO10 and β-actin expression at different time point of RNAi were measured by Western Blot. (E) A549 cells were reverse transfected with siRNAs targeting MYO10 or non-targeting control siRNAs. The transfected cells were re-seeded at 5×10^3^ cells per well into 96 well plates at 48 or 72 hours post transfection (left o right panel, respectively), then infected with RSV-A-GFP 24h later. The infected cells from each time point were moved to Incucyte for live-cell imaging for another 3 days (mean +/- SD, n=3). The area-under-curve (AUC) of percentage green area per image over 72h for each time course was calculated. Statistical significance was determined by two-way ANOVA with multiple comparisons. (**** p < 0.0001).

MYO10 knockout clone 1H5 and A549wt cells were infected with RSV-A-GFP (MOI 0.1), and monitored using Incucyte live-cell imaging. Clone 1H5 exhibited a consistent and significant reduction in RSV infectivity over 72 hours post-infection quantified via GFP fluorescence (Figure 2C).

Given the incomplete validation of a full knockout in clone 1H5, we additionally performed siRNA-mediated knockdown of MYO10 in A549wt cells. Western blot analysis confirmed a robust and sustained reduction of MYO10 protein from 48 to 96 hours post-transfection (Figure 2D). At 72 or 96 hours after siRNA transfection, cells were infected with RSV-A-GFP (MOI 0.1) and monitored for an additional 72 hours by Incucyte live-cell imaging. MYO10 knock down consistently and significantly decreased RSV infectivity (Figure 2E).

In summary, phenotypic validation using both CRISPR-mediated gene disruption and siRNA-mediated knockdown demonstrates that loss of MYO10, partial or complete, significantly reduces RSV infectivity at later stages of infection.

### MYO10 does not modulate IFN induction or signalling but is slightly upregulated by RSV infection

Given previous reports that cytoskeletal remodeling can influence virus sensing and innate immune activation (Acharya, Reis et al. 2022, van Huizen and Gack 2025) we also investigated potential effects of MYO10 on IFN induction and signaling. Interferon response profiling indicated that MYO10 is not a classical interferon-stimulated gene (ISG), as interferon-α treatment did not induce its mRNA expression (Supplementary figure 2A–C, left panels). However, RSV infection modestly increased MYO10 mRNA at later time points in a Jak/STAT-dependent manner, as ruxolitinib blocked this induction (Supplementary figure 2A–C, right panels). Importantly, MYO10 depletion did not measurably alter IFNβ or downstream ISG induction during RSV infection (Supplementary figure 2D–F). Together, these data argue that MYO10 does not function as a canonical antiviral effector in our system.

### Absence of MYO10 impedes cell migration and wound healing

Previous studies have demonstrated the role of MYO10 in regulating cell migration across multiple cell types, including murine melanoblasts (Tokuo, Bhawan et al. 2018), neuronal cells (Yu, Lai et al. 2015), and cancer cell lines such as HeLa and COS7 (Mas, Cristella et al. 2025). Given our observation that MYO10 knockout (KO) and knockdown (KD) in A549 cells significantly reduced RSV infection, we next investigated whether MYO10 influences RSV susceptibility through effects on cell migration. To assess MYO10’s impact on cell migration not only in uninfected cells, but also during RSV infection, we performed scratch assays using MYO10 KO cells (Figure 3A) and siRNA-mediated MYO10 KD cells (Figure 3B), alongside their respective controls. Cells were either mock infected or infected with RSV-A-GFP (MOI 0.1). Twenty-four hours later, scratches were introduced into the monolayers, and wound closure was monitored by Incucyte live-cell imaging. In both KO and KD settings, loss of MYO10 markedly impaired wound healing efficiency compared with wild-type or control cells (Figure 3), consistent with prior reports linking MYO10 to migratory capacity. These data confirm that MYO10-deficincey impairs cell migration and show this applies to cells infected with RSV as well. Therefore, modulating cell migration dynamics or the efficiency of cell-to-cell contact may be a mechanism by which MYO10 affects RSV infection.

**Figure 3.**
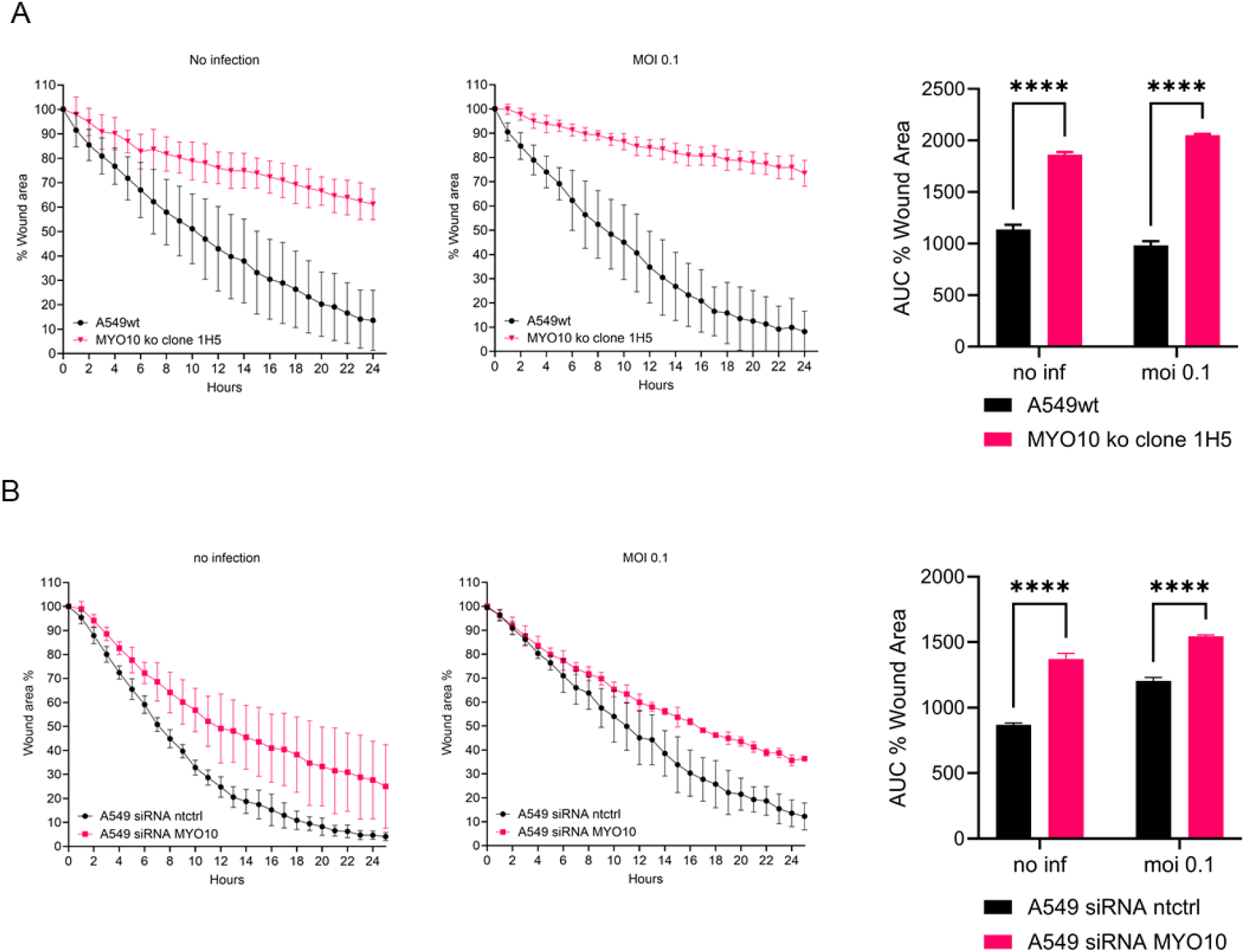
MYO10 knock out and knock down impedes cell migration and wound healing. (A) A549wt and MYO10 knock out clone 1H5 were either mock infected or infected with RSV-A-GFP (MOI 0.1) and 24 hours later scratches were made onto the cell layer and wound healing was monitored through Incucyte live cell imaging. (B) A549 cells were reverse transfected with pool of siRNAs against MYO10 or non-targeting control. Cells were then infected with RSV-A-GFP (MOI 0.1) 48 hours post siRNA transfection and scratches were made one day later and wound healing was monitored through Incucyte live cell imaging. The wound area generated from scratches were quantified via ImageJ plugin Wound-healing-size-tool (https://github.com/AlejandraArnedo/Wound-healing-size-tool/). The data shown are means of three biological replicates. Each replicate represents the mean of at least four scratches. The area-under-curve (AUC) of wound healing curves were calculated. Statistical significance was determined by two-way ANOVA with multiple comparisons. (**** p < 0.0001).

### Loss of MYO10 impairs filopodia formation

MYO10 is well established as a key regulator of filopodia formation (Courson and Cheney 2015). More specifically, over-expression of MYO10 has been shown to strongly upregulate filopodia numbers per cell, while MYO10 silencing decreases endogenous filopodia numbers (Bohil, Robertson et al. 2006, Tokuo, Mabuchi et al. 2007, Watanabe, Tokuo et al. 2010, Courson and Cheney 2015). Filopodia have been implicated in facilitating cell-to-cell spread and intercellular trafficking of viral components for multiple viruses (Schudt, Kolesnikova et al. 2013, Matozo, Kogachi et al. 2022, Zhang, Lin et al. 2024). To investigate how loss of MYO10 alters cell morphology, filopodia formation, and RSV dissemination between neighboring cells, we performed immunofluorescence (IF) staining and confocal microscopy to examine MYO10 localization in the presence or absence of RSV. Cells were stained for MYO10 (green), filamentous actin (F-actin, grey), nucleus (cyan), and, where indicated, RSV F protein (red) (Figure 4).

**Figure 4.**
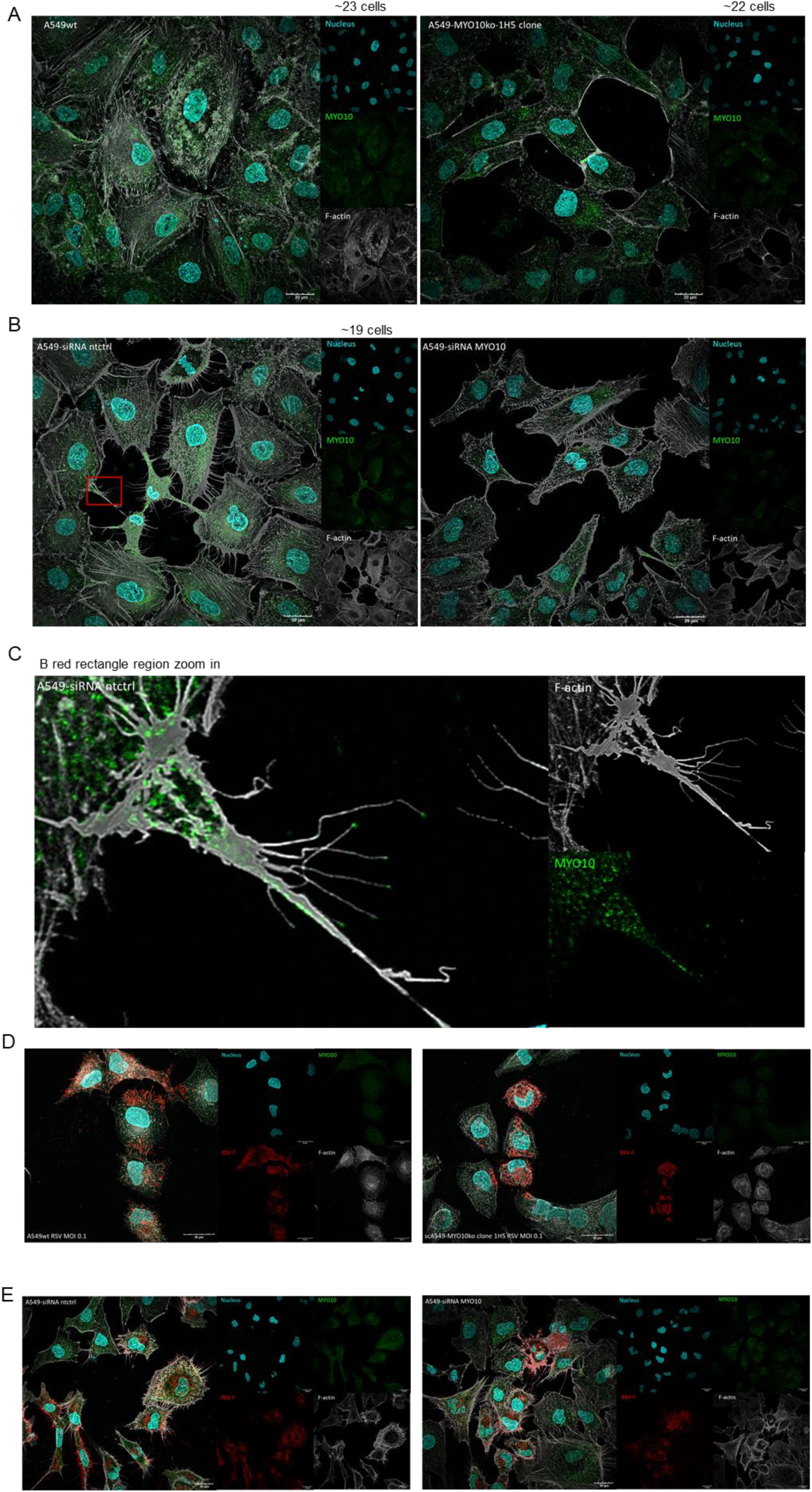
Loss of MYO10 affects filopodia formation and influences RSV cell-to-cell spreading. (A) Confocal immunofluorescence (IF) images of A549wt and MYO10 KO clone 1H5 stained for MYO10 (green), F-actin (gray), and nuclei (DAPI, cyan). Images were acquired using 60x oil objective. Scale bar: 20 µm. A549 cells were reverse transfected with siRNA targeting either MYO10 or non-targeting control for 48h and re-seeded into 24-well plates with glass slips at 1×10^4^ cells/well. One day later, cells were either mock infected (B) or infected with RSV-A-GFP (MOI 0.1) (E) and 48 hours later cells were fixed and stained for RSV-F (red), MYO10 (green), F-actin (gray), and nuclei (DAPI, cyan). Images were acquired using 60x oil objective. Scale bar: 20 µm. (C) Zoomed in image of the red rectangle region in Fig4B of cells reverse transfected with non-targeting control siRNA. (D) Confocal IF images of A549wt and MYO10 KO clone 1H5 infected with RSV-A-GFP (MOI 0.1, 48h post infection) and stained for RSV-F (red), MYO10 (green), F-actin (gray), and nuclei (DAPI, cyan). Images acquired using 100x oil objective. Scale bar: 20 µm. All images displayed are maximum-intensity projections of z-stacks (∼0.2 µm step size) acquired with Olympus FV3000 confocal microscope using 60x or 100x oil objective.

In uninfected cells, loss of MYO10 had a pronounced effect on cellular protrusion formation. Both MYO10 knockout (KO; Figure 4A) and knockdown (KD; Figure 4B) cells exhibited markedly reduced cell-cell contacts compared with control cells, despite similar cell densities within the imaging fields. Consistent with this observation, F-actin staining revealed that actin-driven filopodial structures were shorter and less abundant in MYO10-deficient or -depleted cells. High-magnification imaging occasionally revealed MYO10 localization at filopodial tips (Figure 4C); however, such events were rare, and in most instances MYO10 staining was predominantly cytoplasmic. Thus, in line with previous reports, depletion or ablation of MYO10 expression in A549 cells decreases filopodia formation (Bohil, Robertson et al. 2006, Tokuo, Mabuchi et al. 2007, Watanabe, Tokuo et al. 2010, Courson and Cheney 2015).

### MYO10 KD reduces RSV short- and long-distance spread

Building on our confocal immunofluorescence observations, we next performed an overlay-based spreading assay, conceptually similar to a plaque assay, given that RSV does not form well-defined plaques in A549 cells, to assess the impact of MYO10 depletion on RSV spread to proximal cells when long-distance particle transfer is limited by the viscous overlay (Figure 5). For real-time monitoring using Incucyte live-cell imaging, we selected hydroxypropyl methylcellulose (HPMC) as a transparent viscosity-enhancing agent that is compatible with continuous imaging (Takumi-Tanimukai, Yamamoto et al. 2022).

**Figure 5.**
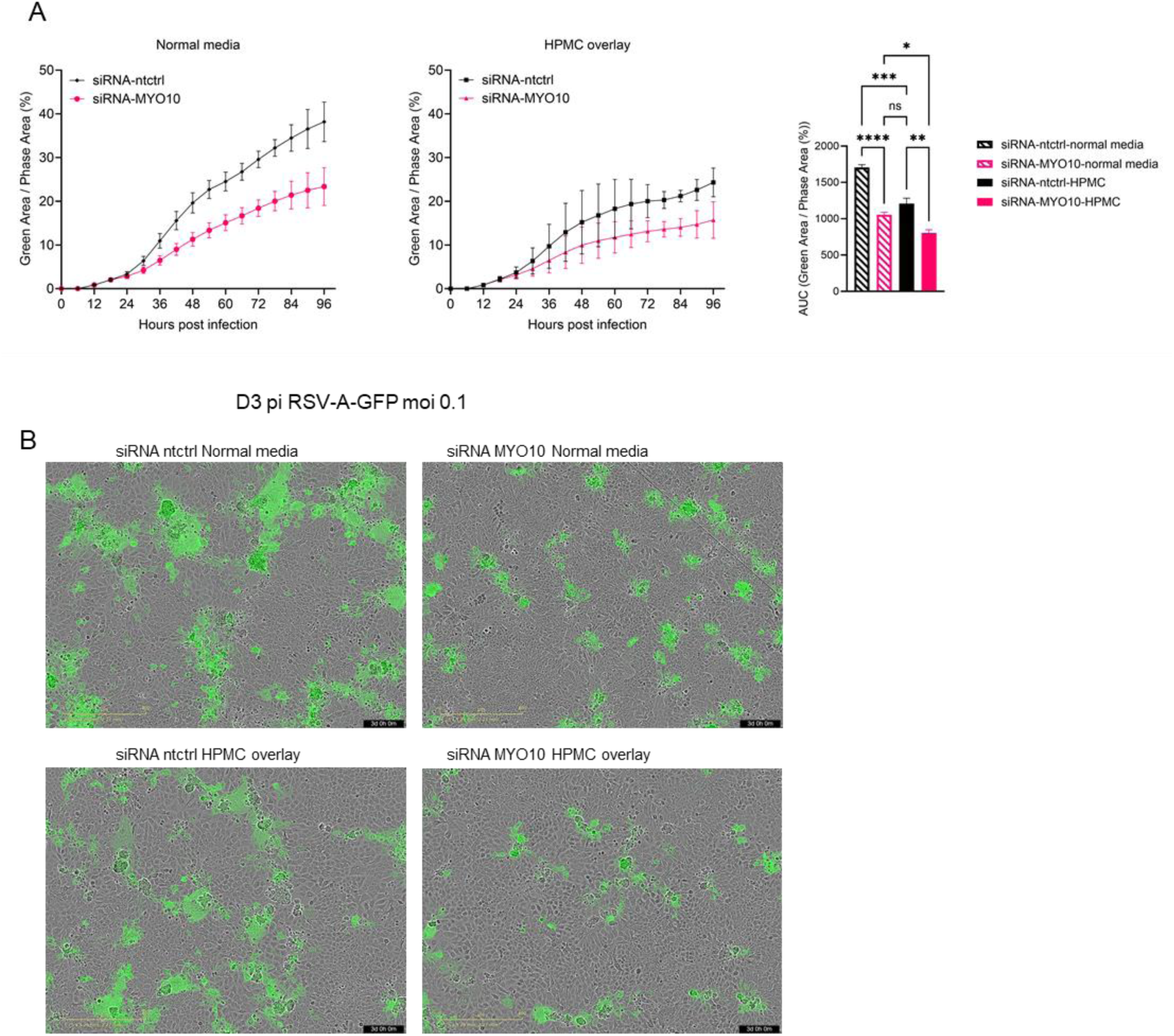
MYO10 KD reduces RSV spreading through culture medium. (A) A549 cells were reverse transfected with siRNA pools targeting MYO10 or non-targeting control for 48 hours then infected with RSV-A-GFP (MOI 0.1) for 4 hours. Virus inoculum was then removed and changed with equal volume of normal media or media containing 0.6% hydroxypropyl methylcellulose (HPMC). Cells were monitored for infectivity using Incucyte live cell imaging, and virus infectivity was measured by normalizing green area to phase area per image per well. Data shown are means of three biological replicates. The area-under-curve (AUC) of infection curves were calculated and statistical analysis was done using one-way ANOVA with multiple comparisons (ns = not significant, *p < 0.05, **p < 0.01, ***p < 0.001, ****p < 0.0001). (B) Representative pictures of cells infected with RSV-A-GFP (MOI 0.1) at 72 hours post infection with or without HPMC in culture media.

Consistent with the expectation that both short distance spread from cell to cell as well as long-distance transfer through the liquid medium contribute to RSV infection dissemination, the presence of HPMC significantly reduced overall RSV spread compared with standard liquid medium (compare Figure 5A and 5B). Importantly, silencing of MYO10 expression reduced the spread of RSV in both, normal media as well as in HPMC overlay culture conditions (Figure 5AB). The effect size of silencing MYO10 was slightly larger in normal media conditions which allows both short and long-distance RSV spread compared to the HPMC overlay, which allows only short distance RSV spread. This is likely because MYO10 directly or indirectly affects both modes of RSV transmission (short and long distance), for instance by directly modulating the efficacy of short distance spread via cell protrusions at cell to cell contact sites and indirectly the number of infected cells that contribute to proximal cell to cell spread and also long-distance transmission through the media. Notably, the early infection kinetics between 0-24h after RSV inoculation revealed no significant difference between the non-targeting control and the MYO10 knock down (Figure 5AB), suggesting that MYO10 expression is dispensable for primary cell entry, RNA transcription and translation.

### MYO10 is important for RSV spread, but not for other steps of the RSV replication cycle

To further test the hypothesis that loss of MYO10 impairs RSV spread and dissemination, we quantified progeny virus production in the culture supernatant (Figure 6). A549 cells were treated with siRNAs, followed 24 hours later by infection with RSV-A-GFP at an MOI of 0.1. Virus-containing supernatants were collected at 48, 72, 96, and 120 hours post infection, and viral release was quantified in parallel by RT-qPCR to measure RSV genome copy numbers (Figure 6A) and by TCID₅₀ assays to determine infectious viral titers (Figure 6B). Although the differences did not reach statistical significance, both assays consistently revealed a trend toward reduced RSV genome copies and lower infectious titers in the supernatant of MYO10 KD cells compared with control cells across all time points examined.

**Figure 6.**
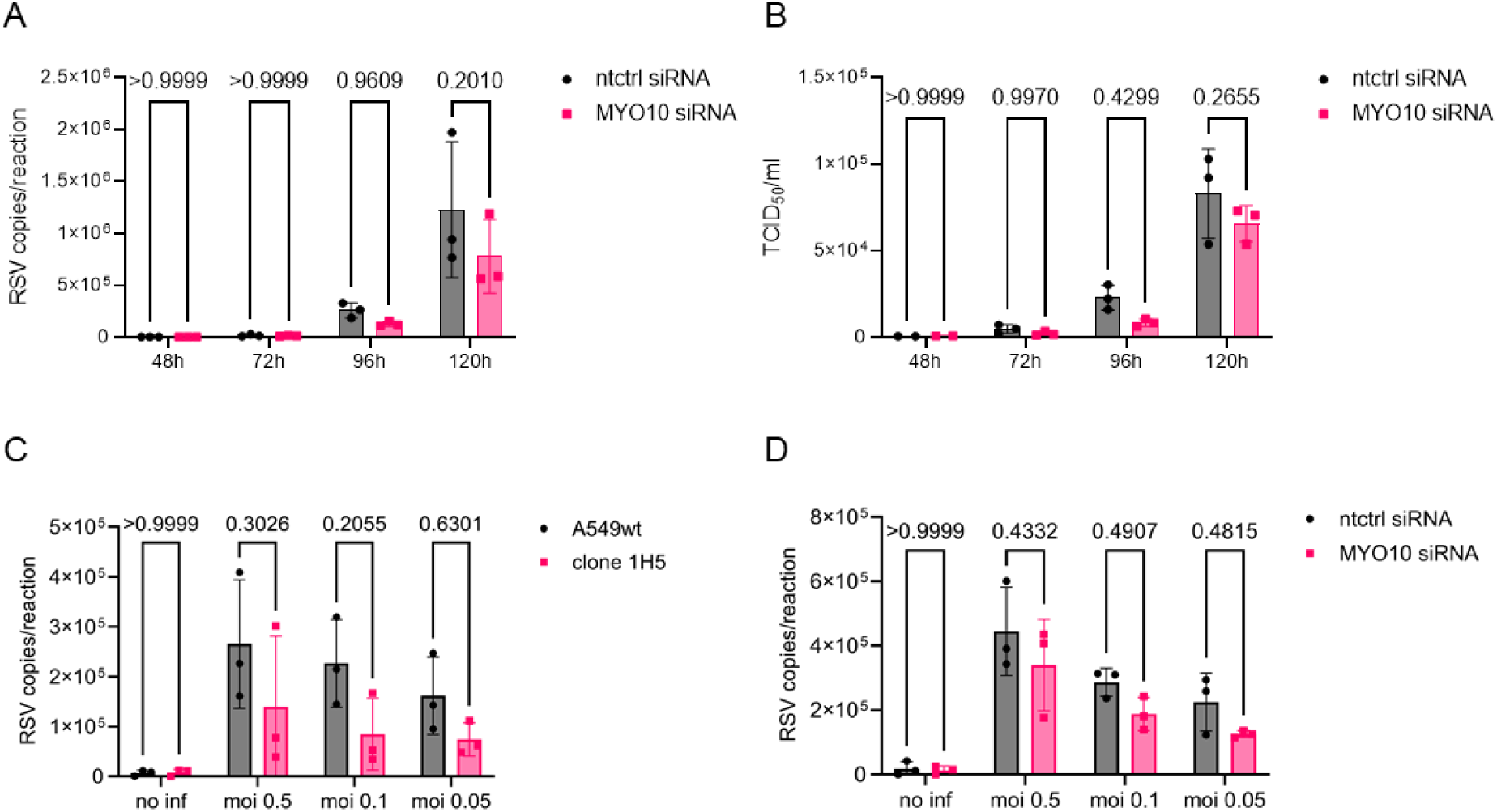
MYO10 KD and KO reduces progeny RSV release. A549 cells were reverse transfected with siRNA targeting either MYO10 or non-targeting control and infected with RSV-A-GFP (MOI 0.1) 24h later. Virus containing supernatants were harvested at 48, 72, 96 and 120 hours post transfection and (A) RSV copy numbers were determined by RT-qPCR (mean +/- SD, n=3). (B) The same virus containing supernatants were used to infect naïve A549 cells and 24 hours later TCID_50_ values were calculated (mean +/- SD, n=3). (C) A549 cells and clone 1H5 were infected with RSV-A-GFP at MOI of 0.5, 0.1, or 0.05 and virus containing supernatants were harvested at 96 hours post infection. RSV copy numbers were determined by RT-qPCR (mean +/- SD, n=3). (D) A549 cells were reverse transfected with siRNA targeting either MYO10 or non-targeting control and infected with RSV-A-GFP (MOI 0.5, 0.1 or 0.05) 24h later. Virus containing supernatants were harvested at 96 hours post transfection and RSV copy numbers were determined by RT-qPCR (mean +/- SD, n=3). Statistical significance in all above assays were calculated via ordinary two-way ANOVA with Sidak’s multiple comparisons test.

To validate this trend using an orthogonal experimental approach, we next quantified RSV genome copies by RT-qPCR in both MYO10 KO clone 1H5 (Figure 6C) and MYO10 KD cells (Figure 6D). Cells were infected with RSV-A-GFP at varying MOIs (0.5, 0.1, and 0.05), and supernatants were harvested at a single time point (96 hours post infection). Consistent with the initial findings, loss of MYO10 resulted in reduced RSV genome copies in the supernatant across all infection conditions tested, although these differences again did not reach statistical significance. These results support the conclusion that MYO10 primarily facilitates the spread of RSV to proximal cells, thereby increasing the number of infected cells. This, in turn, yields a larger quantity of viral progeny that is secreted into the culture fluid.

Previous studies have shown that RSV infection can remodel the composition of the host cell plasma membrane. For example, interaction of the RSV fusion (F) protein with Insulin-like growth factor 1 receptor (IGF1R) promotes the recruitment of nucleolin, an established RSV cellular receptor, to the cell surface, thereby facilitating viral entry (Tayyari, Marchant et al. 2011, Griffiths, Bilawchuk et al. 2020). Given the pronounced role of MYO10 in regulating cell surface architecture and cell-cell interfaces, we wanted to more rigorously test if MYO10 may affect also primary entry and following steps of the RSV replication cycle, prior to spreading to new cells. To address this, we employed palivizumab, a clinically approved human monoclonal antibody targeting RSV F-dependent cell entry, but not other steps of the replication cycle, to terminate RSV infection in a time-dependent manner thereby preventing additional rounds of RSV spread (Figure 7). MYO10 expression was first reduced in A549 cells by reverse transfection with MYO10-targeting siRNA. Two days later, cells were reseeded and infected the following day with RSV-A-GFP at an MOI of 0.1. After a 3-hour virus adsorption period, the inoculum was removed and replaced with either standard culture medium (Figure 7A) or medium containing 8 µg/mL palivizumab (Figure 7B). Infection kinetics were quantified using live-cell imaging and area-under-the-curve (AUC) analysis. As expected, addition of palivizumab strongly inhibited the accumulation of GFP reporter signal over the course of the 72 h follow up (compare Figure 7A and 7B), indicating that a large proportion of GFP expression is due to primary or secondary infection events that occur subsequent to the 4 h inoculation. To specifically assess the efficacy of primary viral entry, transcription, translation and replication, we further quantified infection at 24 hours post inoculation using the Incucyte cell-by-cell analysis module, measuring both the percentage of infected cells (Figure 7D) and the mean GFP fluorescence intensity per cell (Figure 7E). No significant differences were observed between MYO10 KD and control cells in either metric.

**Figure 7.**
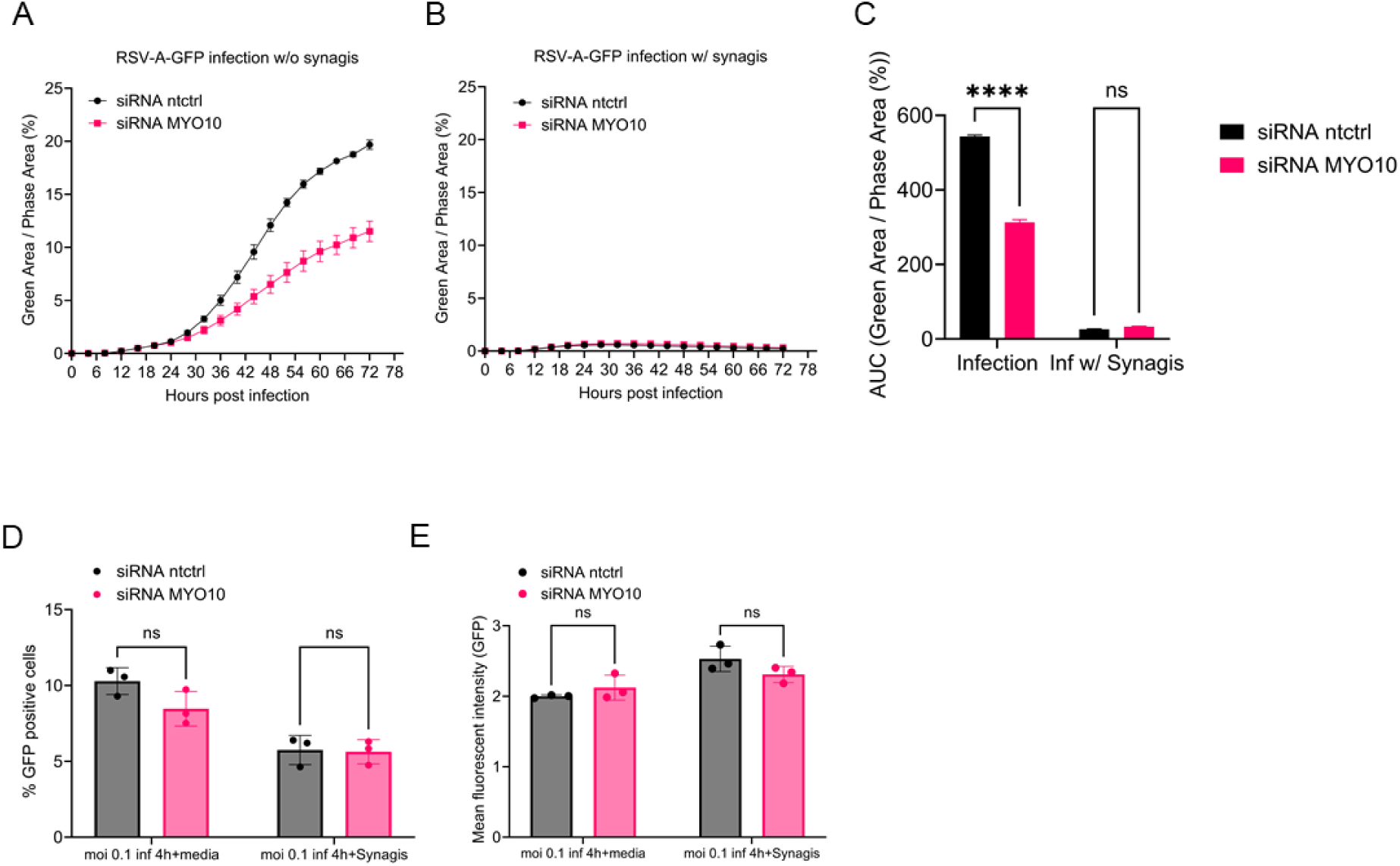
MYO10 knock down does not affect RSV entry. A549 cells were reverse transfected with siRNA targeting MYO10 or non-targeting control for 48 hours, then infected with RSV-A-GFP (MOI 0.05) for 3 hours. Virus inoculum was then removed and changed with (A) fresh media, or (B) media containing 8 µg/mL palivizumab. Cells were then transferred to Incucyte live cell imaging system to monitor infectivity for three days. RSV-A-GFP infectivity was measured via the percentage of green area to phase area per image per well. (C) The area-under-curve (AUC) of infection curves of (A) and (B) were calculated. (D) Percentage of number of infected cells against total cell count was measured at 24h post infection using Incucyte cell-by-cell analysis module. (E) Mean fluorescent intensity of GFP was measured at 24h post infection using Incucyte cell-by-cell analysis module. All statistical analysis were done using two-way ANOVA with multiple comparisons (ns = not significant, *p < 0.05, **p < 0.01, ***p < 0.001, ****p < 0.0001). Data shown are means of three biological replicates.

Together, these results indicate that MYO10 depletion does not impair RSV entry, nor transcription, translation and replication, supporting the conclusion that MYO10 primarily influences viral release and spread.

## Discussion

In this study, we combined human genetics-guided candidate prioritization with functional validation and identified MYO10 as a host factor that modulates RSV infectivity and dissemination. By anchoring our screening strategy in homozygous variants with predicted deleterious effect and enriched in patients with severe RSV disease, we were able to narrow down our search to biologically relevant candidates and ultimately converged on MYO10 as the primary gene whose perturbation consistently attenuated RSV infection in human lung epithelial A549 cells. This integrative approach highlights the value of leveraging patient-derived genetic information to uncover host pathways that shape viral pathogenesis beyond what can be captured by GWAS alone.

The core phenotypes associated with MYO10 loss, including impaired cell migration, defective filopodia formation, and reduced viral spread, are consistent with MYO10’s established role in actin-driven membrane dynamics and protrusion biology. These observations extend previous work implicating actin remodeling and cell protrusions in RSV dissemination (Mehedi, McCarty et al. 2016) and reinforce the notion that filopodia and other actin-based surface architectures can facilitate viral spread in epithelial layers. This concept is supported by broader literature showing that actin-based membrane protrusions, including filopodia, can serve as platforms for virus attachment, directed particle movement (“surfing”), and efficient cell-to-cell transmission across multiple viral families (Mattila and Lappalainen 2008, Chang, Baginski et al. 2016). In our system, MYO10 depletion reduced both long-distance dissemination in standard liquid culture and short-range spread under viscous overlay conditions, consistent with a model in which MYO10-dependent protrusions and cell-cell interfaces increase the efficiency of transmission between neighboring cells and the overall expansion of infected foci.

Our mechanistic experiments further support the conclusion that MYO10 primarily impacts post-entry stages of RSV infection. The absence of early kinetic differences between control and MYO10-depleted cells, combined with palivizumab-based restriction of secondary infection events, argues against a major role for MYO10 in primary entry, early gene expression, or genome replication. Instead, the reduction in extracellular viral genomes and infectious titers, while modest and not always statistically significant, was consistent across independent approaches and time points and aligns with the interpretation that reduced dissemination yields fewer infected producer cells over time, thereby lowering cumulative progeny virus in the supernatant.

Our analysis of MYO10 genetic variation in the IRIS cohort extends these findings toward clinical relevance, but also emphasizes the complexity of translating in vitro phenotypes into disease mechanisms. We identified a homozygous missense variant (rs7737765; H148Y) within the motor domain of MYO10 that is predicted to impair function and occurs in a subset of patients with severe RSV disease (Supplementary figure 3A). While the increased allele frequency of the predicted deleterious minor allele in the IRIS1 cohort was statistically significant at the nominal level (odds ratio of 1.59, 95% CI of 1.06-2.31; Fisher’s exact p = 0.0189), the observed effect size is moderate and the comparison to a reference subpopulation (gnomAD non-Finnish Europeans) rather than a matched control cohort warrants cautious interpretation. Given the limited sample size and potential confounding by population stratification, this association should be considered preliminary pending replication in an independent cohort and functional validation. Functional testing in CRISPR-edited A549 cells revealed a moderate but reproducible reduction in RSV infectivity (Supplementary figure 3B–C). At first glance, reduced infectivity might appear difficult to reconcile with enrichment of this allele in severe RSV cases. However, disease severity *in vivo* is multifactorial and may reflect not only viral replication efficiency but also tissue clearance, epithelial integrity, repair capacity, and immune regulation. Therefore, an MYO10 hypomorphic state could, in principle, produce mixed effects, for example modestly restricting viral dissemination at the cellular level while impairing processes that contribute to effective clearance or recovery of airway function.

One plausible axis how MYO10 loss of function could influence RSV pathogenesis in the lung beyond viral epithelial spread is mucociliary clearance. Efficient elimination of RSV from the airways depends on intact epithelial organization, coordinated repair after injury, and effective mucociliary transport, which is a primary innate defense mechanism of the respiratory tract. Disruption of epithelial architecture or cytoskeletal dynamics can impair mucociliary clearance and prolong pathogen persistence, thereby increasing inflammation and tissue damage (Bustamante-Marin and Ostrowski 2017). Because MYO10 affects cell migration, protrusive structures, and cell–cell connectivity, it is conceivable that MYO10 dysfunction could alter epithelial restitution dynamics or the maintenance of airway surface organization during or after RSV infection. While we have not tested this in differentiated airway cultures, this hypothesis can be addressed in future work using primary human airway epithelium at air–liquid interface and quantitative assays of ciliary beating and mucociliary transport.

A second biologically plausible axis concerns immune cell functions that rely on actin-driven protrusions and membrane remodeling. MYO10 has been implicated in phagocytic cup formation and pseudopod extension in macrophages, acting downstream of PI(3)K signaling; in the original study, inhibition of MYO10 function impaired phagocytosis, including in bovine alveolar macrophages (Cox, Berg et al. 2002). While phagocytosis is not synonymous with immune cell migration, both processes depend on coordinated actin remodeling and membrane protrusion. It is therefore conceivable that MYO10 hypomorphic variants could influence innate immune effector functions in the infected lung, such as particle uptake, clearance of infected debris, or spatial positioning of phagocytes within airway tissue. We emphasize that our study provides no data on MYO10 in immune cells, and any immune-related role remains a hypothesis. Nonetheless, this represents a testable avenue for future investigations using primary macrophages, monocytes, or dendritic cells, ideally combined with tissue-relevant infection models that capture epithelial-immune interactions.

Several limitations should be acknowledged. First, functional experiments were performed in A549 cells, a transformed epithelial cell line that does not fully recapitulate differentiated airway epithelium. Second, viral dissemination and release were quantified in 2D monolayers rather than in organoids or air-liquid interface cultures, precluding direct inference about spread and clearance mechanisms in vivo. Third, the enrichment of rs7737765 in severe RSV cases has not yet been replicated in an independent cohort, and the observed in vitro effect size is modest. Finally, the directionality between cellular phenotypes and clinical severity cannot be inferred from these data alone, underscoring the need for replication genetics and for models that incorporate mucociliary clearance and immune components.

Future studies should therefore validate MYO10’s role in more complex experimental systems and clarify whether MYO10 directly associates with viral components or indirectly shapes a permissive architecture for viral egress and transmission. Expanding genetic analyses to larger cohorts will be essential to validate a potential contribution of MYO10 variants to clinical severity. From a translational perspective, MYO10 itself may not be an immediately tractable therapeutic target; however, our findings highlight the importance of host cytoskeletal pathways in RSV dissemination. Targeting host processes that govern spread, in complement to entry- or replication-directed approaches, may offer strategies to limit disease progression while reducing selective pressure for viral resistance.

In summary, this work connects human genetic variation to a host cytoskeletal regulator that facilitates RSV spread at post-entry stages of infection. By combining patient genomics with functional virology, our study provides new insights into host determinants that influence RSV infection and potentially pathogenesis. We identify MYO10 as a candidate modifier of severe RSV disease, motivating future mechanistic studies in tissue-relevant systems and further clinical validation.

## Materials and Methods

### Study cohort IRIS

This study was conducted on the IRIS cohort comprising 130 children who were hospitalized for severe respiratory illness within the first 36 months of life. Participants were recruited in Germany at different cities. Blood and upper airway samples were collected at the time of admission. Patients were enrolled with confirmed RSV infection via PCR of nasopharyngeal swabs or airway specimens. Exclusion criteria included prematurity, chronic lung disease, congenital heart defects, or immunodeficiencies. In total, 101 individuals were chosen for genetic analysis (Wetzke, Funken et al. 2022).

### Exome sequencing and analysis

Genomic DNA was isolated from whole blood with QIAamp DNA Blood Kit (Qiagen). The exome library was prepared and paired-end sequencing was performed on the Illumina HiSeq 2500 platform with the TruSeq SBS Kit v3, HS. FastQC (v0. 11. 8) (Andrews 2010) was used to assess the sequencing quality. Before mapping to the human reference genome (hg19), reads were trimmed by fastp (v0. 23. 2) (Chen, Zhou et al. 2018) and then aligned using BWA-MEM (Li and Durbin 2009). Picard toolkit’s MarkDuplicates was used to locate duplicate DNA reads and mark them (Wysoker A 2013). The marked BAM files were then used for variant calling following the Genome Analysis Toolkit (GATK) Best Practices workflow. First, variants were called per sample using HaplotypeCaller in GVCF mode (Poplin, Ruano-Rubio et al. 2018) followed by joint genotyping across all samples using GenotypeCaller. Single nucleotide variants (SNVs) and short insertions and deletions were further processed through quality score recalibration using VariantRecalibrator (Auwera and O’Connor 2020).

Each variant was annotated with its allele frequency in the gnomAD Non-Finnish European (NFE) population, zygosity, coding effect (synonymous or nonsynonymous), and genomic location. For variant level annotation, we applied CADD (Rentzsch, Witten et al. 2019), Condel (González-Pérez and López-Bigas 2011), and PROVEAN (Choi and Chan 2015) while for gene-level constraint, we used the LOEUF20 score (Karczewski, Francioli et al. 2020). Variant annotations were conducted through SnpEff (Cingolani, Platts et al. 2012).

### Selection of interferon (IFN)-related genes

A set of 5,141 IFN-related genes was collected from the Interferome database (Rusinova, Forster et al. 2013) an in-house collection of interferon-regulated genes in primary human cells (Lauber, Kazem et al. 2015), and Ingenuity Pathway Analysis (IPA). IPA analysis targeted interferon signalling, pathogen-sensing pathways, and RSV-related functions. Homozygous variants located within these IFN-related genes were prioritized for candidate filtering.

### Candidate filtering

Homozygous single-nucleotide variants (SNVs) were first extracted from annotated exome sequencing data and defined as Level 0 candidates. This initial dataset comprised more than 41,000 SNPs identified in 101 patients with severe RSV infection, mapping to 2,299 genes.

Subsequently, two parallel filtering strategies based on minor allele frequency (MAF) were applied. In the first approach, variants with a gnomAD MAF < 0.05 were retained. In the second approach, variants showing a difference in MAF (MAF_diff) greater than 0.05 between the IRIS cohort and the gnomAD reference population were selected. Together, these two filtering branches reduced the candidate set to approximately 400 genes, collectively designated as Level 1 candidates.

To further prioritize potentially deleterious variants, in silico variant effect prediction tools (CADD, Condel, and PROVEAN) and a gene-level constraint metric (loss-of-function observed/expected upper bound fraction; LOEUF) were applied. Overlap between the different filtering criteria was visualized using a Venn diagram, and genes harboring variants predicted to be deleterious by at least two prediction tools were prioritized. This refined set of candidates was defined as Level 2.

Using this integrative filtering strategy, a final set of 23 genes containing high-confidence candidate SNPs was identified for downstream functional validation. Detailed information on these genes and variants is provided in Supplementary table S1.

### Cells, viruses and media

A549 (ATCC CCL-185), HEp-2 (ATCC CCL-23), and HEK293T (ATCC CCL-3216) cells were cultured in F-12K Nutrient Mixture, Advanced MEM, or DMEM, respectively, supplemented with 10% heat-inactivated fetal calf serum (FCS; Capricorn Scientific), 1% non-essential amino acids (NEAA; Gibco), 2 mM L-glutamine (Gibco), and 100 U/ml penicillin and 100 µg/ml streptomycin (Gibco). Media with the aforementioned supplements are denoted as complete medium throughout this paper. Cells were maintained at 37 °C in a humidified incubator with 5% CO₂. Recombinant reporter virus rHRSV-A-GFP (Rameix-Welti, Le Goffic et al. 2014) and clinical isolate RSV-A-ON1-H1 (Sake, Zhang et al. 2024) have been described previously.

### CRISPR/Cas9 knock out screening of candidate genes

Single-guide RNA (sgRNA) sequences targeting candidate genes (Supplementary table S3) were designed using the CCTop tool (Stemmer, Thumberger et al. 2015, Labuhn, Adams et al. 2018) and cloned into the SGL40C.EFS.dTomato (SGL40C.EFS.dTomato was a gift from Dirk Heckl (Addgene plasmid # 89395; http://n2t.net/addgene:89395 ; RRID:Addgene_89395)) (Reimer, Knöß et al. 2017) vector via BsmBI restriction enzyme-mediated ligation of phosphorylated oligonucleotides. The resulting sgRNA-expressing plasmids were co-transfected with the envelope plasmid pCZ.VSV-G (Kalajzic, Stover et al. 2001) and packaging plasmid pCMV.ΔR8.74 (Dull, Zufferey et al. 1998) into HEK293T cells to produce lentiviral particles. A549 cells were transduced with a lentiviral vector carrying a Cas9 expression cassette and selected using 5 µg/ml blasticidin (Heckl, Kowalczyk et al. 2014). pLKO5d.EFS.SpCas9.P2A.BSD was a gift from Benjamin Ebert (Addgene plasmid # 57821; http://n2t.net/addgene:57821; RRID:Addgene 57821). Lentiviruses were harvested and used to transduce selected A549/Cas9 cells. CRISPR knockouts cells targeting each candidate gene were generated and, 72 hours post-transduction, infected with rHRSV-A-GFP at a multiplicity of infection (MOI) of 0.1 for 48 hours. RSV infectivity was assessed by measuring GFP fluorescence while sgRNA transduction efficiency was controlled by dTomato fluorescence via flow cytometry.

### CRISPR RNP transfection and generation of single cell clones

In vitro generation of sgRNAs targeting MYO10 were conducted according to the protocol from NEB (NEB #E3322V/S).

Oligo nucleotide synthesized for sgRNA generation:

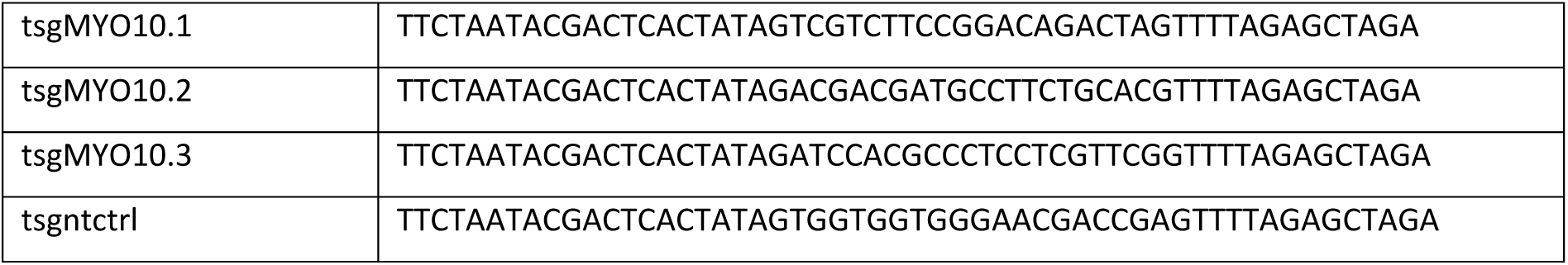

Expected sgRNA sequence after synthesis:

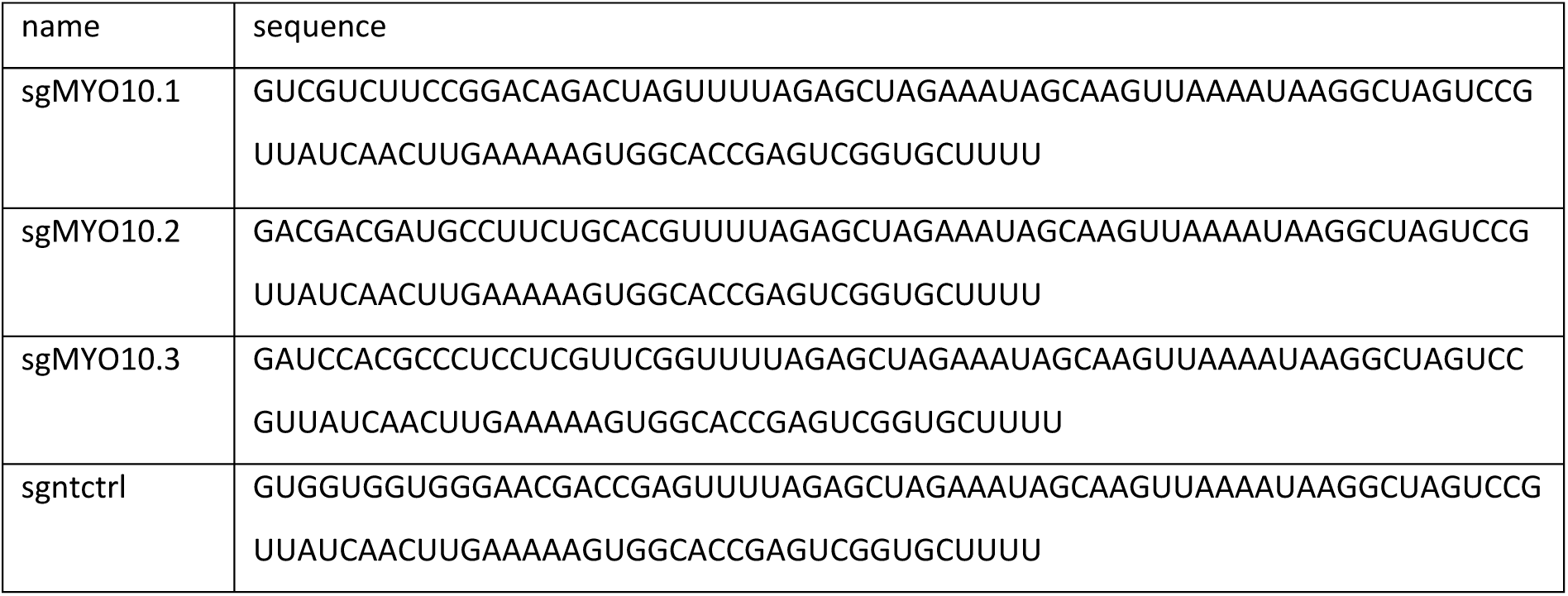

After generation of sgRNA, RNP complex containing each sgRNA and Cas9 protein was generated according to the NEB protocol (Transfection of Cas9 RNP (ribonucleoprotein) into adherent cells using the Lipofectamine® RNAiMAX). A549 cells were reverse transfected together with the RNP complex via Lipofectamine RNAiMAX, and 48 hours post transfection, cells were seeded for single cell selection in 96 well plates in the presence of ROCK inhibitor Y-27632 (10µM) to boost cell survival and single clone growth. Single colonies were observed in the following days and moved to larger format for further culturing. The knockout efficiency of the single cell clones were validated by Sanger sequencing and indel frequency analysis using the TIDE tool (Brinkman, Chen et al. 2014), then further confirmed via Western Blot.

### siRNA knock down of MYO10

MYO10 knockdown was performed via reverse transfection using a pool of four small interfering RNAs (siRNAs) targeting MYO10 or a non-targeting control (Dharmacon ON-TARGETplus siRNA SMARTpool). siRNAs were dissolved in RNase-free 1× siRNA buffer (Dharmacon) at a concentration of 20 µM for storage at –20 °C. For transfection, siRNAs were diluted to a working concentration of 5 µM in RNase-free H₂O and mixed with Opti-MEM at a 1:60 (v/v) ratio. A second master mix containing Lipofectamine RNAiMAX at 1:150 (v/v) in Opti-MEM was prepared and combined with the siRNA mixture in equal volume. For reverse transfection, 300 µl of the siRNA–lipid complex was mixed with 300 µl of A549 cell suspension (2.5 × 10⁴ cells) and seeded into one well of a 24-well plate. After 24 hours, the medium was replaced with fresh complete media, and cells were incubated at 37 °C with 5% CO₂ for 48 to 120 hours before further analysis.

siRNA sequence:

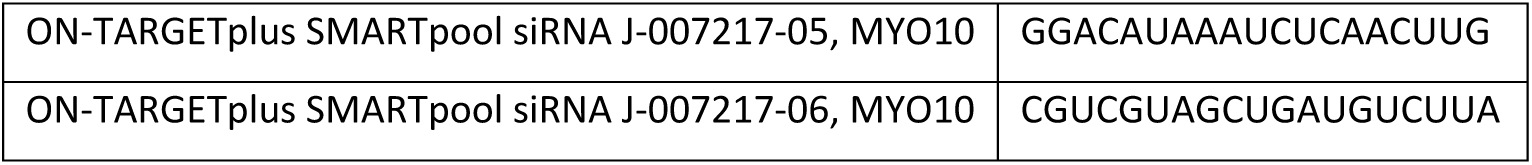
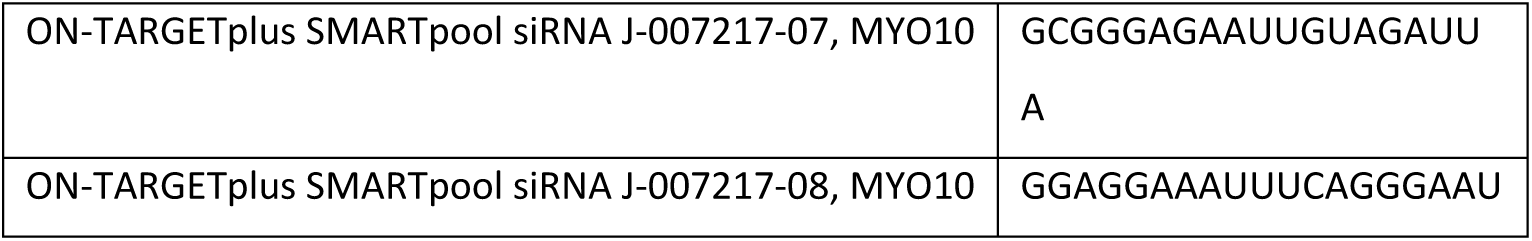

### Live-cell imaging with Incucyte

Cells were seeded into 96- or 24-well plates at the indicated densities and subjected to the specified treatments. Plates were then transferred to the Incucyte SX5 live-cell imaging system (Sartorius) and equilibrated for 30 minutes prior to the first image acquisition. Unless otherwise specified, live-cell imaging throughout this study was performed using the adherent cell-by-cell analysis mode with a 10× objective and the Green/Orange/Near-Infrared (NIR) optical module, applying default exposure settings. Images were acquired at defined intervals and durations according to each experimental setup. Image processing and quantitative analyses were conducted using either the Incucyte Basic Analysis module or the Incucyte Cell-by-Cell Analysis module. Output metrics, including cell confluence (%) and total fluorescence area/Total integrated intensity (GCU/OCU/NIRCU × µm² per image), were subsequently exported and analyzed using GraphPad Prism software.

### Immunofluorescence imaging

Cells were seeded onto glass coverslips placed in 24-well plates and infected the following day with RSV-A-H1^32^ at a MOI of 0.1. After 48 hours, cells were fixed with 3% paraformaldehyde (PFA), permeabilized with 0.5% Triton X-100 in PBS, and subsequently immunostained. Primary antibodies included anti-MYO10 (rabbit polyclonal, Atlas Antibodies; Cat# HPA024223; RRID: AB_1854248) and anti-RSV fusion protein (Palivizumab, a human monoclonal antibody targeting RSV-A F protein; AbbVie Ltd, North Chicago, IL). Secondary antibodies were goat anti-rabbit IgG AlexaFluor 488 (ThermoFisher) and goat anti-human IgG AlexaFluor 568 (ThermoFisher). F-actin was visualized using phalloidin-iFluor 647 (Abcam; Cat# ab176759), and nuclei were stained with DAPI. Confocal imaging was performed using an FV3000 microscope (Olympus) with 60× or 100× objectives. Acquired images were processed by constrained iterative deconvolution and maximum intensity projection.

### Scratch assay

Cells were seeded in 24-well plates at a density of 5 × 10⁴ cells per well in 500 µL of complete media. The following day, cells were infected with RSV-A-GFP at a MOI of 0.1, with uninfected cells serving as controls. At 24 hours post-infection, linear scratches were introduced using 100 µL pipette tips.

Wells were gently washed with PBS to remove detached cells and replenished with fresh complete media. Plates were then transferred to the Incucyte live-cell imaging system, and wound closure was monitored by time-lapse imaging at 1-hour intervals over a 24-hour period. Images were analyzed using ImageJ, and wound areas were quantified using the Wound Healing Size Tool plugin (Suarez-Arnedo, Torres Figueroa et al. 2020). Quantified wound areas were subsequently plotted using GraphPad Prism software.

### Plaque assay with hydroxypropyl methylcellulose (HPMC)

Cells were seeded into 24-well plates at a density of 2.5 × 10⁴ cells per well in 500 µL of complete culture medium. After 24 hours, cells were infected with RSV-A-GFP at a MOI of 0.1 for 3 hours. Following infection, the virus inoculum was removed and replaced with either complete F12K nutrient mixture medium or medium containing HPMC. The HPMC overlay was prepared by combining ¼ volume of 2x MEM (pH adjusted with 1/10 volume of sodium bicarbonate), ¼ volume of 2.4% (w/v) HPMC (dissolved in H_2_O and sterilized by autoclaving), and ½ volume of F12K nutrient mixture medium supplemented with 1% penicillin/streptomycin and 2% FCS. The final overlay medium contained 0.6% (w/v) HPMC. Following overlay application, plates were transferred to the Incucyte live-cell imaging system, and infection loci was monitored by time-lapse imaging at 6-hour intervals over a period of 4 days. Quantitative data were exported and analyzed using GraphPad Prism.

### Western blot

Cells were washed once with PBS, then detached and lysed in RIPA buffer (1% (v/v) Triton X-100, 2mM EDTA (pH 8), 100mM Tris-HCl (pH 7.4), 300mM NaCl) with protein inhibitor (10 ml RIPA + 1 tablet of protease inhibitor (Roche #04693124001)). Lysates were centrifuged at 1,000 × g for 10 minutes to remove cellular debris. The resulting supernatant (total cell lysate) was collected, and protein concentration was determined using the Bradford assay. For sodium dodecyl sulfate–polyacrylamide gel electrophoresis (SDS-PAGE), 20 µg of total protein was mixed with 2x SDS sample buffer (100mM Tris-HCl, 4% (w/v) SDS, 20% (v/v) glycerol, 200mM DTT, 0.0018% (w/v) bromophenol blue), heated at 95 °C for 8 minutes, and loaded onto an 8% separation gel. Electrophoresis was performed at 110 V for 2 hours, followed by transfer of proteins onto PVDF membranes using the Trans-Blot Turbo system (Bio-Rad). Membranes were blocked in 5% (w/v) non-fat milk in PBST (0.5% v/v Tween-20 in PBS), then incubated overnight at 4 °C with antibodies against MYO10 (Atlas Antibodies; rabbit polyclonal; Cat# HPA024223; RRID: AB_1854248) or β-actin (Thermo Fisher Scientific; β-Actin Monoclonal Antibody [BA3R], HRP-conjugated; Cat# MA5-15739-HRP). The following day, membranes were washed five times with PBST. Membranes probed for MYO10 were subsequently incubated with an HRP-conjugated goat anti-rabbit IgG secondary antibody. Detection was performed using chemiluminescence on the Vilber Fusion FX imaging system.

### Reverse transcription and real-time quantitative PCR (RT-qPCR)

Cells from one well of a 24-well plate were lysed in 350 µl RA1 buffer (NucleoSpin RNA Kit, Macherey-Nagel) supplemented with 1:100 (v/v) β-mercaptoethanol, and lysates were stored at −20 °C until further processing. Total RNA was extracted using the NucleoSpin RNA Kit according to the manufacturer’s instructions. RNA concentration was measured using a NanoDrop spectrophotometer (Thermo Fisher Scientific), and samples were stored at −80 °C. For each RT-qPCR reaction, 250 ng of total RNA was reverse-transcribed into cDNA using the PrimeScript Reverse Transcription Kit (Takara) following the manufacturer’s protocol. The resulting cDNA was used as a template for quantitative PCR (qPCR) with TB Green Premix Ex Taq II (Takara) and gene-specific primer pairs. qPCR was performed on a LightCycler 96 system (Roche), and relative gene expression levels were calculated using the 2^−ΔΔCT^ method (Livak and Schmittgen 2001).

RT-qPCR primer pairs:

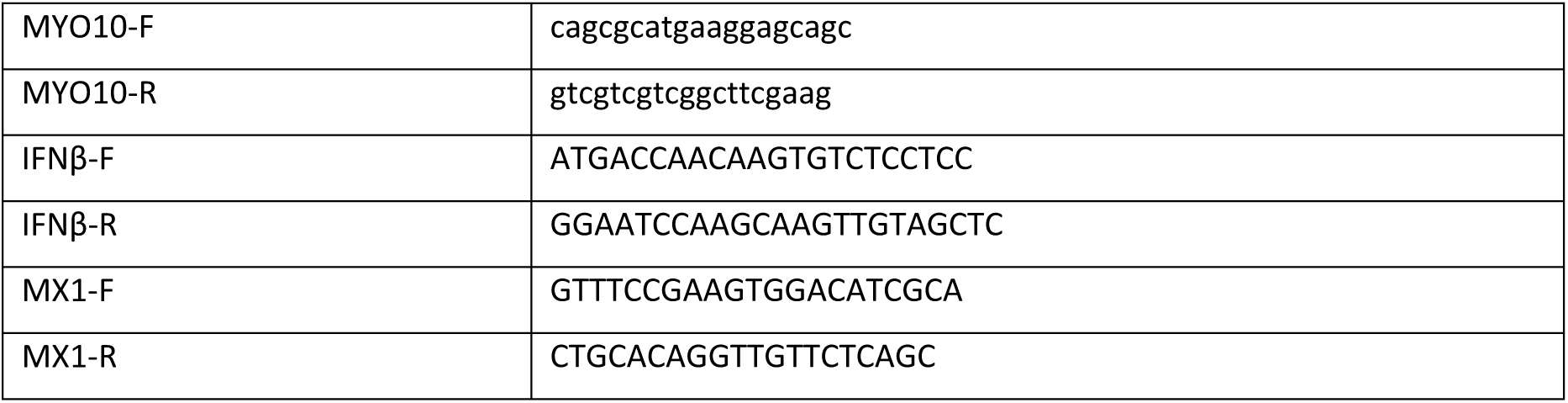

### RSV copy number determination by RT-qPCR

RSV-containing cell culture supernatants were collected into 1.5 mL microcentrifuge tubes and centrifuged to remove cellular debris. Cleared supernatants were aliquoted (200 µL per aliquot) and either processed immediately or stored at −70 °C until use. Viral RNA was extracted from supernatants using the High Pure Viral RNA Kit (Roche; cat. no. 11858882001) according to the manufacturer’s instructions. RNA was eluted in 50 µL of nuclease-free water and stored at −70 °C prior to analysis. RSV genome copy numbers were quantified using the Luna® Probe One-Step RT-qPCR 4× Mix with UDG (New England Biolabs; M3029) following the manufacturer’s protocol, with an RSV-specific probe. A standard curve was generated using a pcDNA3 plasmid containing a partial RSV F gene sequence. Ten-fold serial dilutions of the pcDNA3-RSV-F plasmid (10⁹ to 10¹ copies), with nuclease-free water as a no-template control, were included in each run. RT-qPCR was performed on a LightCycler 96 system (Roche), and RSV genome copy numbers in samples were calculated based on the standard curve.

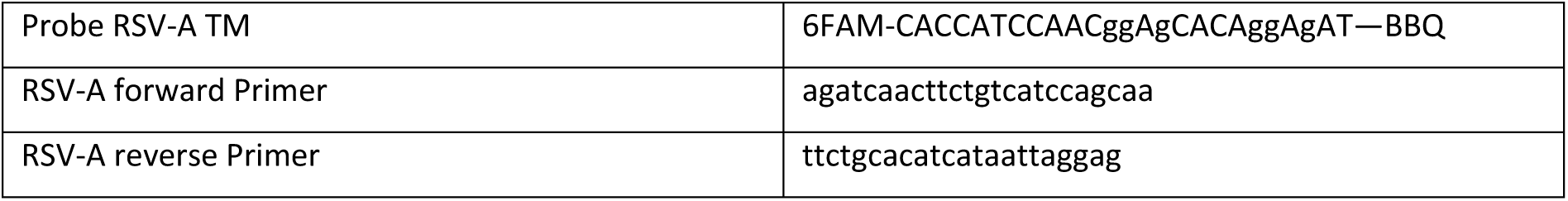

### CRISPR editing

A549 cells with MYO10 H148Y (rs7737765) variant were generated by electroporation with CRISPR-Cas9 RNP (18 pmol Alt-R Cas9 and 22 pmol Alt-R crRNA:tracrRNA duplex) together with 100 pmol of HDR donor DNA oligonucleotide (sequences in Supplementary table S4). 5×10^5^ cells were electroporated using the Neon transfection system (1,600 V, 10 ms, 3 pulses; 10 µl tips) and then transferred to pre-warmed medium supplemented with 1 µM HDR enhancer V2. Medium was changed after 24 h and after another 72 h single cell clones were plated by limiting dilution. Single-cell clones with homo- or heterozygous H148Y alleles were identified by Sanger sequencing (Microsynth, Göttingen, Germany) of the PCR-amplified genomic locus. Electropherograms were analyzed using Benchling (https://www.benchling.com/).

## Acknowledgements

I.N., C.L. and T.P. were funded by the Deutsche Forschungsgemeinschaft (DFG, German Research Foundation) under Germany’s Excellence Strategy - EXC 2155 - project number 390874280. TP received funding from the Volkswagen Stiftung within the Niedersächsische Vorab initiative and the INDIRA: INtegrative Data analytics for Respiratory syncytial virus risk Assessment project. T.Z. was supported by collaborative research center TRR237 (DFG project number 369799452).

Portions of this manuscript were edited for clarity and language with the assistance of a generative artificial intelligence–based language model. The AI tool was used to support writing and revision and did not generate original data, analyses, or scientific conclusions. All content was critically reviewed, verified, and approved by the authors, who take full responsibility for the integrity and accuracy of the work.

We thank Marie-Anne Rameix-Welti and Jean-François Eléouët for the kind gift of the rHRSV-A-GFP virus. We are grateful to Birthe Reinecke for project management support and all members of the Institute for Experimental Virology for advice and constructive discussion of this study.

---

**Supplementary Figure 1.**
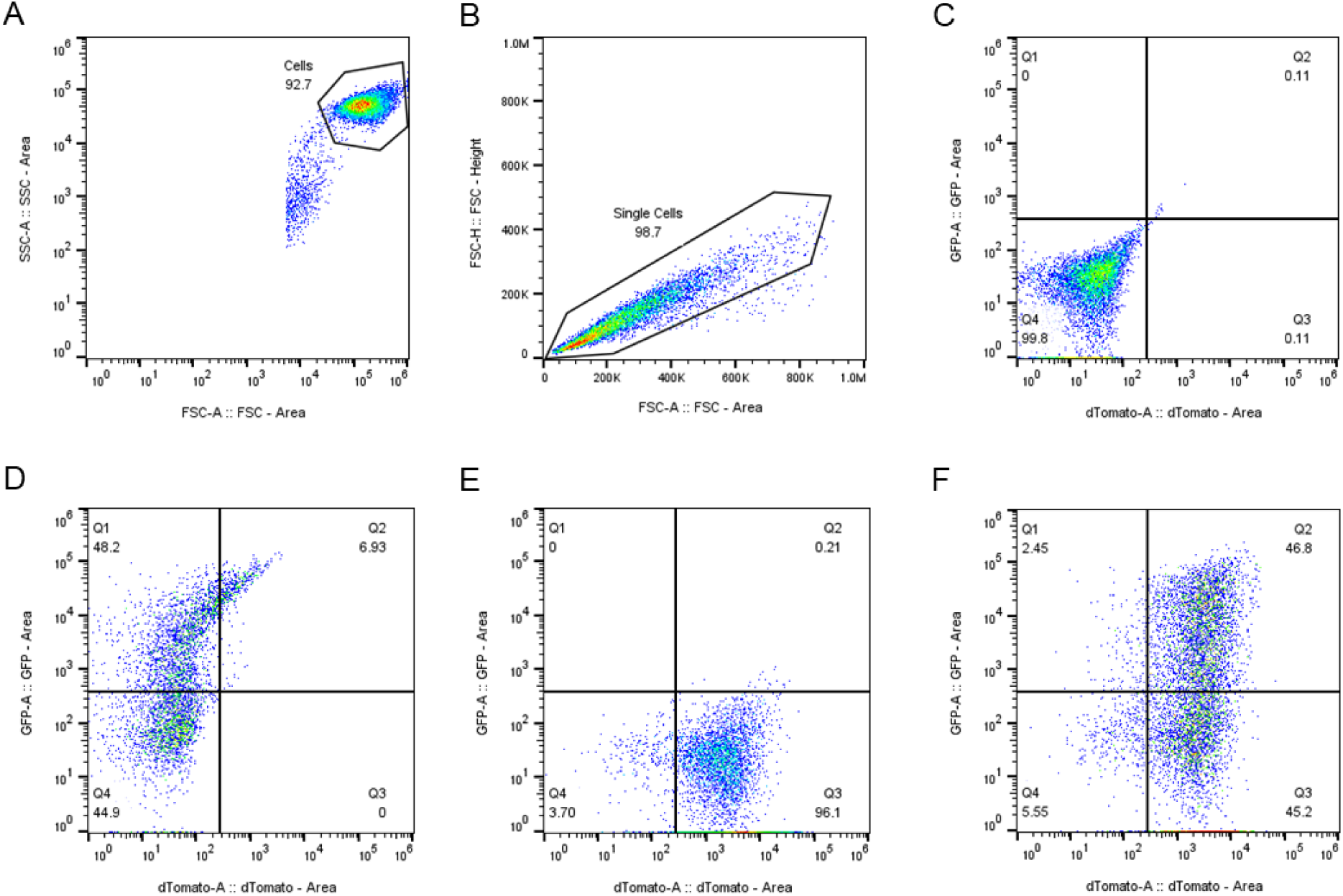
Representative gating strategy. (A) FSC-A vs SSC-A to select cells and exclude debris; (B) FSC-A vs FSC-H for selecting single cells and exclude doublets/triplets; (C) marker gating for non-transfected and non-infected cells to identify double negative population; (D) marker gating for cells infected with RSV-A-GFP to identify GFP+ and GFP– cells; (E) marker gating for cells transduced with lentiviral particles with dTomato reporter to identify dTomato+ and dTomato– cells; (F) gating for cells transduced with lentiviral particles with dTomato marker and infected with RSV-A-GFP. Double positive cell population is depicted in Q2.

**Supplementary Figure 2.**
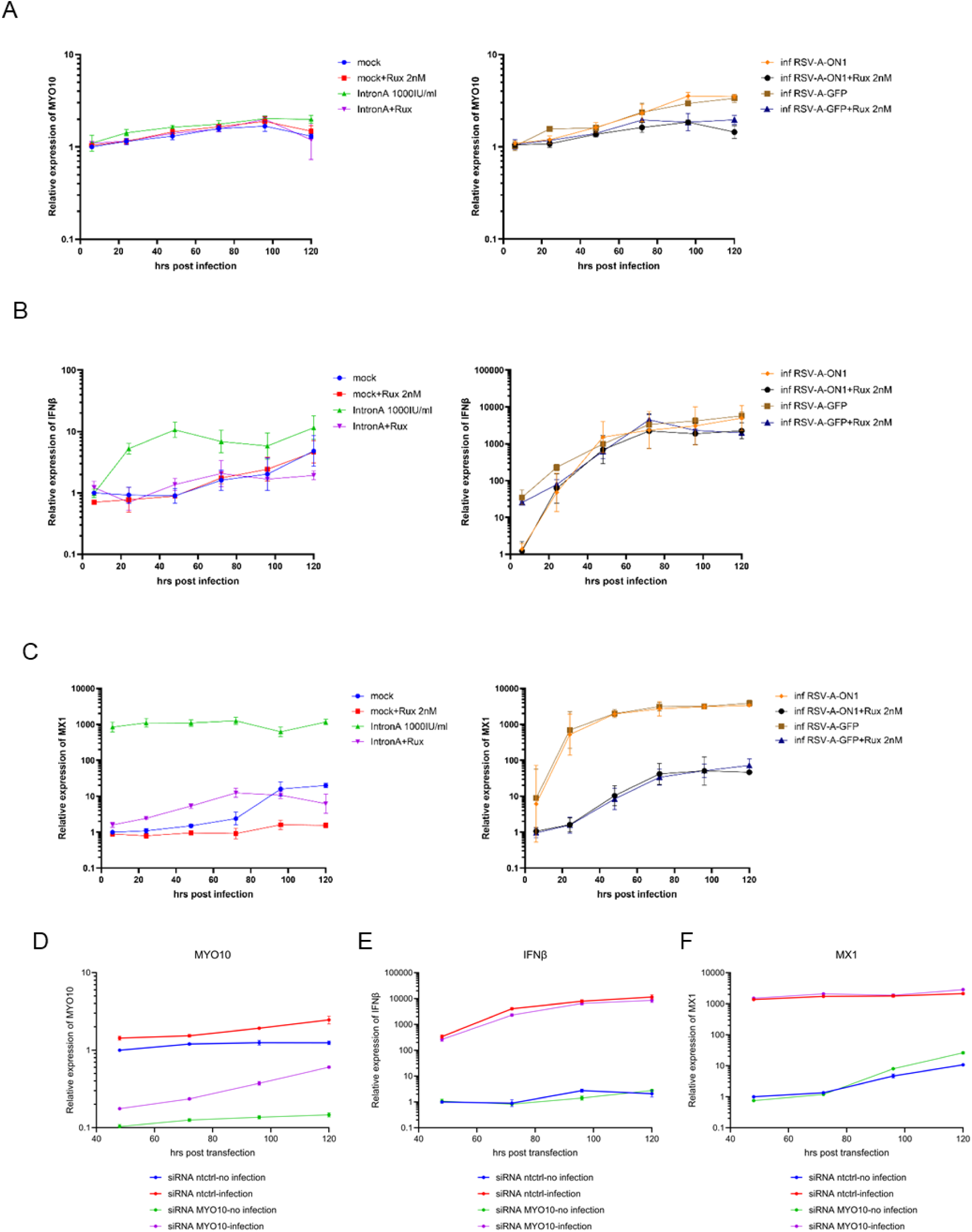
MYO10 expression is slightly induced by RSV infection and has no impact on IFN induction and downstream signaling. (A, B, C) A549 cells were seeded at 1×10^4^ cells/well in 24-well plates and were treated with either DMSO (mock) or ruxolitinib (2 nM). One day later, the cells were treated with DMSO or 1000 IU/ml of intron A or infected with RSV-A-GFP or RSV-A-H1 with MOI of 0.05. After 4 hours virus inoculum was removed, and cells were replenished with fresh media either with or without ruxolitinib (2 nM). Total intracellular RNA was extracted at 6, 24, 48, 72, 96, 120 hours post infection or treatment. The mRNA level for (A) MYO10, (B) IFNβ, and (C) MX1 were quantified via RT-qPCR (mean +/- SD, n=3). (D, E, F) A549 cells were reverse transfected with siRNA targeting MYO10 or non-targeting control for 48 hours, then infected with RSV-A-GFP (MOI 0.05) for 4 hours or incubated with fresh media for 4 hours (no infection). Virus inoculum or cell culture fluid was then removed and changed with fresh media. Total intracellular RNA was extracted at 48, 72, 96, 120 hours post inoculation. The mRNA level for (D) MYO10, (E) IFNβ, and (F) MX1 were quantified via RT-qPCR (mean +/- SD, n=3).

**Supplementary Figure 3.**
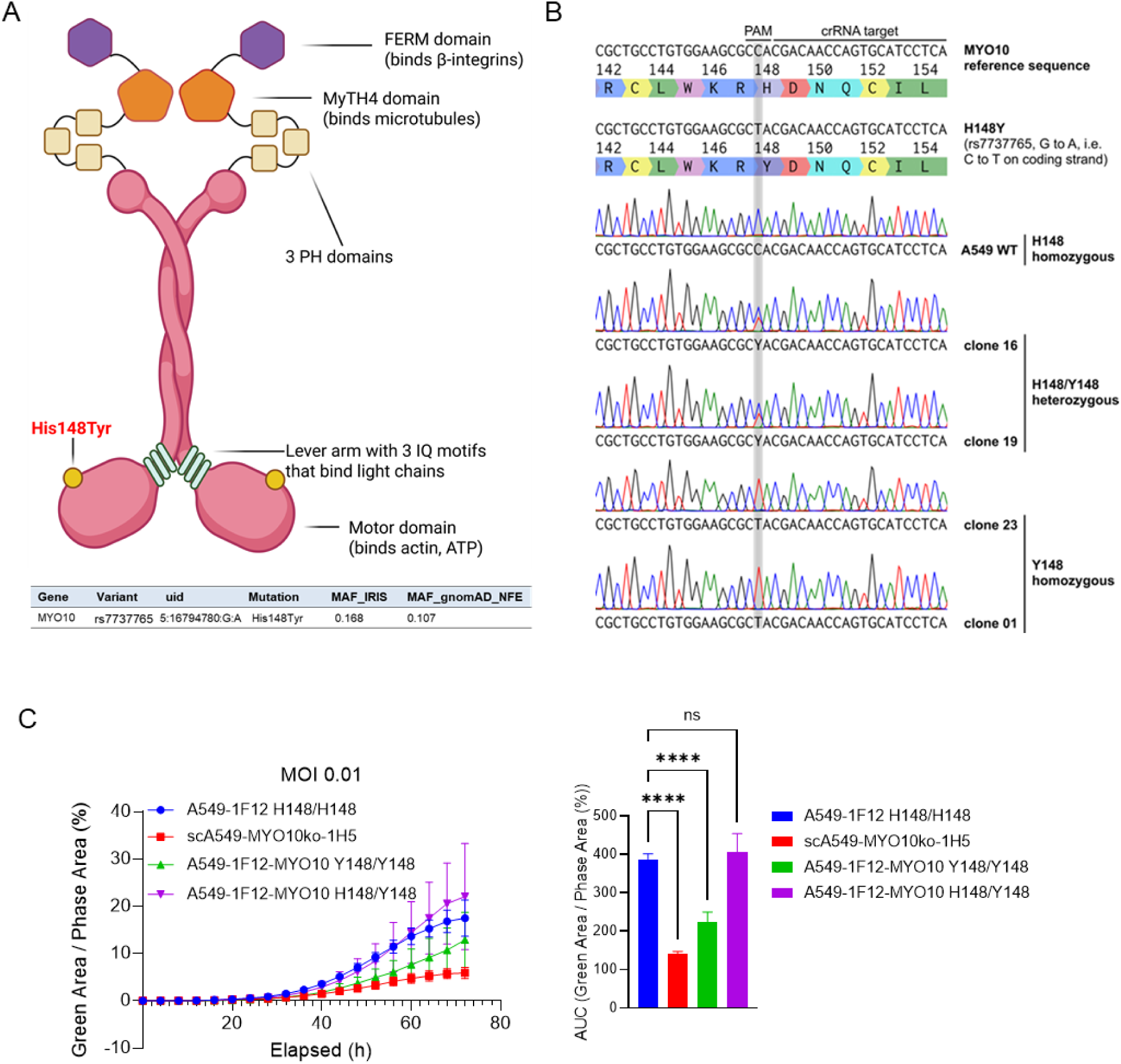
MYO10 mutation His148Tyr influences RSV infectivity. (A) Schematic illustration of MYO10 structure domain and the relative location of MYO10 Histidine 148. The SNP rs7737765 results in a tyrosine mutation at position 148, which is over-represented in the IRIS cohort. (B) Single nucleotide edition was performed on A549wt-1F12 clone via CRISPR knock in and two clones for either heterozygous or homozygous were selected. (C) Edited cell pools together with parental cell clone and one MYO10 KO clone were infected with RSV-GFP reporter virus and tested via live-cell imaging (mean +/- SD, n=3). The area under- curve (AUC) of percentage green area per image over 72h was calculated. (Two-way ANOVA, ** = p < 0.01, *** = p < 0.001, **** = p < 0.0001).

**Supplementary Table S1.** List of candidate risk genes and observed SNVs. uid = chromosome:position:referenceAllele:alternativeAllele; type: UTR = 5’ or 3’ untranslated region, NSM = non-synonymous missense, NSN = non-synonymous nonsense, NSF = non-synonymous frameshift; MAF_hRSV = alternative allele frequency in the n101 hRSV cohort; MAF_gnomAD_NFE = alternative allele frequency in gnomAD non-finish Europeans; MAF_diff = MAF difference between hRSV and gnomAD; RAF_gnomAD_NFE = reference allele frequency in gnomAD non-finish Europeans; CADD =scaled (phred) CADD score relative to all possible variants; LOEUF = LOEUF score decile bin; CONDEL = Condel score; PROVEAN_PREDICTION = PROVEAN prediction (cutoff < -2.5); NA = not available.

| Gene name | Variant ID | uid | type | MAF_hRSV | MAF_gnomAD_NFE | MAF_diff | RAF_gnomAD_NFE | CADD | LOEUF | CONDEL | PROVEAN_PREDICTION |
| --- | --- | --- | --- | --- | --- | --- | --- | --- | --- | --- | --- |
| ACAN | rs1042630 | 15:89402051:A:G | NSM | 0,782 | 0,748171 | -0,033829 | 0,251829 | 0,002 | 1 | 0,328347475 | Neutral |
| ACAN | rs1126823 | 15:89417238:A:G | NSM | 0,215 | 0,236431 | -0,021431 | 0,763569 | 19,5 | 1 | 0,39339909 | Neutral |
| ACAN | rs2882676 | 15:89400339:A:C | NSM | 0,728 | 0,644137 | -0,083863 | 0,355863 | 12,88 | 1 | 0,52124431 | Neutral |
| ACAN | rs34949187 | 15:89386652:G:A | NSM | 0,203 | 0,173894 | 0,029106 | 0,826106 | 28 | 1 | 0,366241622 | Deleterious |
| ACAN | rs3743398 | 15:89398407:C:T | NSM | 0,282 | 0,206577 | 0,075423 | 0,793423 | 23,5 | 1 | 0,328886853 | Deleterious |
| ACAN | rs3817428 | 15:89415247:C:G | NSM | 0,332 | 0,271615 | 0,060385 | 0,728385 | 23 | 1 | 0,375108254 | Deleterious |
| ACAN | rs4932439 | 15:89401109:A:G | NSM | 0,822 | 0,823731 | 0,001731 | 0,176269 | 11,05 | 1 | 0,215477997 | Neutral |
| ACAN | rs938608 | 15:89398605:G:T | NSM | 0,728 | 0,642835 | -0,085165 | 0,357165 | 15,85 | 1 | 0,29189998 | Deleterious |
| ACAN | rs938609 | 15:89398631:T:A | NSM | 0,723 | 0,642347 | -0,080653 | 0,357653 | 16,42 | 1 | 0,5781153 | Neutral |
| AHNAK | rs11231128 | 11:62290719:A:G | NSM | 0,03 | 0,031045 | -0,001045 | 0,968955 | 16,12 | 2 | 0,125699609 | Neutral |
| AHNAK | rs61312994 | 11:62285441:A:G | NSM | 0,03 | 0,031016 | -0,001016 | 0,968984 | 22,5 | 2 | 0,263284624 | Neutral |
| AHNAK | rs74853209 | 11:62296601:T:C | NSM | 0,03 | 0,030998 | -0,000998 | 0,969002 | 19,23 | 2 | 0,58311463 | Neutral |
| AHNAK | rs75436331 | 11:62297447:T:C | NSM | 0,03 | 0,030945 | -0,000945 | 0,969055 | 16,06 | 2 | 0,37327178 | Neutral |
| AHNAK | rs75855515 | 11:62297162:G:A | NSM | 0,025 | 0,030951 | -0,005951 | 0,969049 | 15,41 | 2 | 0,294188132 | Neutral |
| AHNAK | rs77055528 | 11:62296535:T:C | NSM | 0,03 | 0,030995 | -0,000995 | 0,969005 | 22,1 | 2 | 0,305414274 | Deleterious |
| ANKS1B | rs11109968 | 12:100042040:C:G | NSM | 0,307 | 0,308998 | -0,001998 | 0,690993 | 17,93 | 1 | 0,374830778 | NA |
| ANKS1B | rs11109969 | 12:100042163:A:G | NSM | 0,307 | 0,309248 | -0,002248 | 0,690752 | 7,599 | 1 | 0,330688727 | NA |
| ANKS1B | rs77436242 | 12:99640159:G:C | NSM | 0,074 | 0,048941 | 0,025059 | 0,951059 | 23 | 1 | 0,472826366 | Neutral |
| ANKS1B | rs77436242 | 12:99640159:G:C | NSM | 0,074 | 0,048941 | 0,025059 | 0,951059 | 23 | 1 | 0,527777778 | Neutral |
| AP2A2 | rs11246379 | 11:1009307:C:T | UTR | 0,094 | 0,082377 | 0,011623 | 0,917623 | 0,027 | 1 | NA | NA |
| AP2A2 | rs144441591 | 11:988619:A:G | NSM | 0,02 | 0,004016 | 0,015984 | 0,995984 | 24,6 | 1 | 0,559965541 | Deleterious |
| AP2A2 | rs61869001 | 11:1011597:G:A | NSM | 0,158 | 0,114878 | 0,043122 | 0,885122 | 10,12 | 1 | NA | NA |
| AP2A2 | rs72842410 | 11:1010649:C:T | NSM | 0,144 | 0,13554 | 0,00846 | 0,86446 | 4,26 | 1 | 0,316388467 | Neutral |
| AP2A2 | rs7950955 | 11:1011490:G:A | NSM | 0,55 | 0,544648 | -0,005352 | 0,455352 | 4,119 | 1 | NA | NA |
| ATG2A | rs35115827 | 11:64679362:G:A | NSM | 0,059 | 0,043929 | 0,015071 | 0,956071 | 20,2 | 2 | 0,35068151 | Neutral |
| ATG2A | rs35115827 | 11:64679362:G:A | NSM | 0,059 | 0,043929 | 0,015071 | 0,956071 | 20,2 | 2 | 0,458829365 | Neutral |
| ATG2A | rs60711419 | 11:64679331:G:A | NSM | 0,059 | 0,043921 | 0,015079 | 0,956079 | 18,95 | 2 | 0,354339848 | Neutral |
| ATG2A | rs61746813 | 11:64677273:C:T | NSM | 0,009901 | 0,000349 | 0,009552 | 0,999651 | 22,7 | 2 | 0,340744932 | Neutral |
| ATG2A | rs61746813 | 11:64677273:C:T | NSM | 0,009901 | 0,000349 | 0,009552 | 0,999651 | 22,7 | 2 | 0,624503968 | Neutral |
| CCND3 | rs1051130 | 6:41903782:A:C | NSM | 0,579 | 0,545776 | -0,033224 | 0,454224 | 9,205 | 1 | 0,405064302 | Neutral |
| CCND3 | rs1051130 | 6:41903782:A:C | NSM | 0,579 | 0,545776 | -0,033224 | 0,454224 | 9,205 | 1 | 0,430912281 | Neutral |
| CCND3 | rs11552783 | 6:42016319:G:A | UTR | 0,188 | 0,186603 | 0,001397 | 0,813397 | 11,69 | 1 | NA | NA |
| CCND3 | rs33966734 | 6:41903798:C:A | NSM | 0,015 | 0,013587 | 0,001413 | 0,986413 | 21 | 1 | 0,393302993 | NA |
| CCND3 | rs33966734 | 6:41903798:C:A | NSM | 0,015 | 0,013587 | 0,001413 | 0,986413 | 21 | 1 | 0,393302993 | Neutral |
| CCND3 | rs33966734 | 6:41903798:C:A | NSM | 0,015 | 0,013587 | 0,001413 | 0,986413 | 21 | 1 | 0,627480159 | NA |
| CCND3 | rs33966734 | 6:41903798:C:A | NSM | 0,015 | 0,013587 | 0,001413 | 0,986413 | 21 | 1 | 0,627480159 | Neutral |
| CENPE | rs1381657 | 4:104061993:C:G | NSM | 0,213 | 0,181931 | 0,031069 | 0,818069 | 0,325 | 2 | 0,46785378 | Neutral |
| CENPE | rs143398214 | 4:104115535:C:T | NSM | 0,009901 | 0,000412 | 0,009489 | 0,999588 | 26,6 | 2 | 0,528199891 | Deleterious |
| CENPE | rs2243682 | 4:104059542:G:A | NSM | 0,158 | 0,208534 | -0,050534 | 0,791466 | 11,49 | 2 | 0,526090628 | Neutral |
| CENPE | rs2615542 | 4:104066461:A:G | NSM | 0,163 | 0,211176 | -0,048176 | 0,788824 | 1,376 | 2 | 0,363114376 | Neutral |
| DMXL2 | rs12102203 | 15:51791559:A:G | NSM | 0,475 | 0,520249 | 0,045249 | 0,479741 | 11,31 | 1 | 0,405044042 | Neutral |
| DMXL2 | rs143033674 | 15:51791234:G:A | NSM | 0,009901 | 0,003049 | 0,006852 | 0,996951 | 28,7 | 1 | 0,448695677 | Deleterious |
| DMXL2 | rs17524906 | 15:51829812:G:A | NSM | 0,252 | 0,237554 | 0,014446 | 0,762446 | 13,1 | 1 | 0,375272607 | Neutral |
| DSP | rs2076299 | 6:7580958:A:G | NSM | 0,074 | 0,041712 | 0,032288 | 0,958288 | 17,2 | 1 | 0,470424525 | Neutral |
| DSP | rs34884895 | 6:7585795:C:A | NSM | 0,009901 | 0,000019 | 0,009882 | 0,999981 | 22,7 | 1 | 0,528353391 | Deleterious |
| DSTYK | rs35845538 | 1:205138321:G:C | NSM | 0,045 | 0,049731 | -0,004731 | 0,950269 | 22 | 2 | 0,584553503 | Neutral |
| DSTYK | rs3851294 | 1:205130413:A:G | NSM | 0,901 | 0,904313 | 0,003313 | 0,095687 | 22,9 | 2 | 0,395497873 | Neutral |
| E2F5 | rs12926 | 8:86089787:C:G | UTR | 0,615 | 0,718475 | 0,103475 | 0,281486 | 11,92 | 2 | NA | NA |
| E2F5 | rs143705255 | 8:86127239:T:A | UTR | 0,009901 | 0,000196 | 0,009705 | 0,999804 | 20,8 | 2 | 0,59672619 | NA |
| FAM135A | rs2747701 | 6:71238105:A:G | NSM | 0,416 | 0,471094 | -0,055094 | 0,528906 | 22,5 | 1 | 0,422403352 | Deleterious |
| MINK1 | rs11556634 | 17:4796274:T:C | NSM | 0,416 | 0,36931 | 0,04669 | 0,630676 | 19,69 | 1 | 0,292878787 | Neutral |
| MINK1 | rs11556635 | 17:4796286:C:T | NSM | 0,406 | 0,348251 | 0,057749 | 0,651749 | 22,9 | 1 | 0,349637861 | Neutral |
| MINK1 | rs2302319 | 17:4797305:G:A | NSM | 0,089 | 0,075114 | 0,013886 | 0,924886 | 19,07 | 1 | 0,592698696 | Neutral |
| MINK1 | rs79940062 | 17:4797910:G:A | NSM | 0,025 | 0,019855 | 0,005145 | 0,980145 | 26,9 | 1 | 0,562105109 | Neutral |
| MYC | rs4645959 | 8:128750540:A:G | NSM | 0,05 | 0,034067 | 0,015933 | 0,965933 | 25,2 | 1 | 0,420969181 | Deleterious |
| MYO10 | rs11750538 | 5:16769273:G:A | NSM | 0,584 | 0,600761 | 0,016761 | 0,399239 | 23,8 | 2 | 0,52746122 | Deleterious |
| MYO10 | rs11750538 | 5:16769273:G:A | NSM | 0,584 | 0,600761 | 0,016761 | 0,399239 | 23,8 | 2 | 0,52746122 | Neutral |
| MYO10 | rs149110 | 5:16701717:C:T | NSM | 0,025 | 0,005645 | 0,019355 | 0,994355 | 10,73 | 2 | NA | NA |
| MYO10 | rs17707947 | 5:16877744:C:T | NSM | 0,02 | 0,008747 | 0,011253 | 0,991253 | 14,69 | 2 | 0,527274089 | Neutral |
| MYO10 | rs25901 | 5:16673975:C:G | NSM | 0,624 | 0,665326 | 0,041326 | 0,334644 | 21,3 | 2 | 0,457368142 | Neutral |
| MYO10 | rs7737765 | 5:16794780:G:A | NSM | 0,168 | 0,115096 | 0,052904 | 0,884904 | 23 | 2 | 0,57310267 | Deleterious |
| PLEC | rs11136334 | 8:145001588:C:T | NSM | 0,401 | 0,418548 | -0,017548 | 0,581452 | 17,29 | 2 | 0,369400455 | Neutral |
| PLEC | rs11136336 | 8:145007187:G:A | NSM | 0,381 | 0,39466 | -0,01366 | 0,60534 | 17,08 | 2 | 0,428709051 | Neutral |
| PLEC | rs12543539 | 8:145020477:T:C | UTR | 0,381 | 0,401342 | -0,020342 | 0,598658 | 5,551 | 2 | NA | NA |
| PLEC | rs138924815 | 8:145009036:C:T | NSM | 0,02 | 0,018747 | 0,001253 | 0,981253 | 27 | 2 | 0,51379195 | Deleterious |
| PLEC | rs138924815 | 8:145009036:C:T | NSM | 0,02 | 0,018747 | 0,001253 | 0,981253 | 27 | 2 | 0,51379195 | Neutral |
| PLEC | rs35027700 | 8:144994916:C:T | NSM | 0,02 | 0,001948 | 0,018052 | 0,998052 | 10,55 | 2 | 0,49709487 | Neutral |
| PLEC | rs55895668 | 8:145001031:T:C | NSM | 0,436 | 0,425667 | 0,010333 | 0,574333 | 16,13 | 2 | 0,266865079 | Neutral |
| PLEC | rs55895668 | 8:145001031:T:C | NSM | 0,436 | 0,425667 | 0,010333 | 0,574333 | 16,13 | 2 | 0,288468848 | Neutral |
| PLEC | rs7002002 | 8:144997927:G:A | NSM | 0,401 | 0,42345 | -0,02245 | 0,57655 | 15,22 | 2 | 0,52846663 | Neutral |
| PLEC | rs7833924 | 8:144996029:A:G | NSM | 0,436 | 0,42352 | 0,01248 | 0,57648 | 15,99 | 2 | 0,443031058 | Neutral |
| POU2F1 | rs77105959 | 1:167341211:C:T | NSM | 0,009901 | 0 | 0,009901 | 1 | 26,4 | 2 | 0,541753018 | NA |
| POU2F1 | rs77105959 | 1:167341211:C:T | NSM | 0,009901 | 0 | 0,009901 | 1 | 26,4 | 2 | 0,541753018 | Neutral |
| STRN3 | rs2273171 | 14:31381351:T:C | NSM | 0,401 | 0,46754 | -0,06654 | 0,53246 | 22,4 | 2 | 0,477700832 | Deleterious |
| STRN3 | rs2281563 | 14:31495346:C:T | UTR | 0,125 | 0,113441 | 0,011559 | 0,886559 | 21,8 | 2 | 0,49735791 | Neutral |
| TNFAIP3 | rs2230926 | 6:138196066:T:G | NSM | 0,035 | 0,033431 | 0,001569 | 0,966569 | 20,1 | 1 | 0,425148308 | Deleterious |
| TNS1 | rs11680854 | 2:218747073:A:C | NSM | 0,02 | 0,019091 | 0,000909 | 0,980909 | 23,3 | 2 | 0,58203417 | Deleterious |
| TNS1 | rs11680854 | 2:218747073:A:C | NSM | 0,02 | 0,019091 | 0,000909 | 0,980909 | 23,3 | 2 | 0,58203417 | Neutral |
| TNS1 | rs11680854 | 2:218747073:A:C | NSM | 0,02 | 0,019091 | 0,000909 | 0,980909 | 23,3 | 2 | 0,673611111 | Deleterious |
| TNS1 | rs11680854 | 2:218747073:A:C | NSM | 0,02 | 0,019091 | 0,000909 | 0,980909 | 23,3 | 2 | 0,673611111 | Neutral |
| TNS1 | rs2571445 | 2:218683154:A:G | NSM | 0,584 | 0,609481 | 0,025481 | 0,390519 | 16,85 | 2 | 0,367133812 | Neutral |
| TNS1 | rs3796028 | 2:218695102:G:A | NSM | 0,381 | 0,347703 | 0,033297 | 0,652297 | 12,58 | 2 | 0,472312299 | Neutral |
| TNS1 | rs3796033 | 2:218713282:G:A | NSM | 0,332 | 0,311589 | 0,020411 | 0,688411 | 19,39 | 2 | 0,542996657 | Neutral |
| TNS1 | rs7600493 | 2:218770003:G:A | UTR | 0,416 | 0,375594 | 0,040406 | 0,624406 | 2,442 | 2 | NA | NA |
| TNS1 | rs918949 | 2:218674697:C:T | NSM | 0,579 | 0,603999 | 0,024999 | 0,395991 | 7,39 | 2 | 0,309644589 | Neutral |
| TTN | rs1001238 | 2:179464527:T:C | NSM | 0,272 | 0,228234 | 0,043766 | 0,771766 | 23,3 | 2 | 0,380747923 | Neutral |
| TTN | rs1001238 | 2:179464527:T:C | NSM | 0,272 | 0,228234 | 0,043766 | 0,771766 | 23,3 | 2 | NA | Neutral |
| TTN | rs10497520 | 2:179644855:T:C | UTR | 0,861 | 0,867845 | 0,006845 | 0,132155 | 22,4 | 2 | 0,47783105 | Deleterious |
| TTN | rs10497520 | 2:179644855:T:C | UTR | 0,861 | 0,867845 | 0,006845 | 0,132155 | 22,4 | 2 | 0,47783105 | Neutral |
| TTN | rs12463674 | 2:179432185:A:G | NSM | 0,243 | 0,304741 | -0,061741 | 0,695259 | 23,4 | 2 | 0,504808813 | Deleterious |
| TTN | rs12463674 | 2:179432185:A:G | NSM | 0,243 | 0,304741 | -0,061741 | 0,695259 | 23,4 | 2 | 0,593755193 | Deleterious |
| TTN | rs12463674 | 2:179432185:A:G | NSM | 0,243 | 0,304741 | -0,061741 | 0,695259 | 23,4 | 2 | NA | Deleterious |
| TTN | rs12693164 | 2:179579093:T:C | NSM | 0,178 | 0,150565 | 0,027435 | 0,849435 | 13,48 | 2 | 0,498773612 | Neutral |
| TTN | rs12693166 | 2:179587130:C:G | NSM | 0,178 | 0,150359 | 0,027641 | 0,849641 | 23,5 | 2 | 0,508281987 | Deleterious |
| TTN | rs13021201 | 2:179449131:G:A | NSM | 0,035 | 0,039995 | -0,004995 | 0,960005 | 24,4 | 2 | 0,501378695 | Deleterious |
| TTN | rs13021201 | 2:179449131:G:A | NSM | 0,035 | 0,039995 | -0,004995 | 0,960005 | 24,4 | 2 | NA | Deleterious |
| TTN | rs13390491 | 2:179582327:C:T | NSM | 0,178 | 0,149972 | 0,028028 | 0,850028 | 21,1 | 2 | 0,534561054 | Neutral |
| TTN | rs16866378 | 2:179393111:A:G | NSM | 0,015 | 0,020668 | -0,005668 | 0,979332 | 13,94 | 2 | 0,507371666 | Neutral |
| TTN | rs16866378 | 2:179393111:A:G | NSM | 0,015 | 0,020668 | -0,005668 | 0,979332 | 13,94 | 2 | 0,507421605 | Neutral |
| TTN | rs16866378 | 2:179393111:A:G | NSM | 0,015 | 0,020668 | -0,005668 | 0,979332 | 13,94 | 2 | NA | Neutral |
| TTN | rs16866406 | 2:179457147:G:A | NSM | 0,198 | 0,14446 | 0,05354 | 0,85554 | 23 | 2 | 0,523687912 | Deleterious |
| TTN | rs16866406 | 2:179457147:G:A | NSM | 0,198 | 0,14446 | 0,05354 | 0,85554 | 23 | 2 | NA | Deleterious |
| TTN | rs16866465 | 2:179583496:T:G | NSM | 0,178 | 0,150434 | 0,027566 | 0,849566 | 19,84 | 2 | 0,474896675 | Neutral |
| TTN | rs16866538 | 2:179659912:G:A | NSM | 0,064 | 0,051877 | 0,012123 | 0,948123 | 16,75 | 2 | 0,531793723 | Deleterious |
| TTN | rs2042995 | 2:179558366:T:C | NSM | 0,267 | 0,227651 | 0,039349 | 0,772349 | 21,4 | 2 | 0,357764011 | Neutral |
| TTN | rs2042995 | 2:179558366:T:C | NSM | 0,267 | 0,227651 | 0,039349 | 0,772349 | 21,4 | 2 | NA | Neutral |
| TTN | rs2042996 | 2:179451420:G:A | NSM | 0,272 | 0,224395 | 0,047605 | 0,775605 | 23,2 | 2 | 0,227330808 | Neutral |
| TTN | rs2042996 | 2:179451420:G:A | NSM | 0,272 | 0,224395 | 0,047605 | 0,775605 | 23,2 | 2 | NA | Neutral |
| TTN | rs2163008 | 2:179528378:C:T | NSM | 0,069 | 0,059645 | 0,009355 | 0,940355 | 18,21 | 2 | NA | NA |
| TTN | rs2244492 | 2:179554305:C:T | NSM | 0,317 | 0,3568 | -0,0398 | 0,6432 | 23,2 | 2 | 0,416166726 | Neutral |
| TTN | rs2244492 | 2:179554305:C:T | NSM | 0,317 | 0,3568 | -0,0398 | 0,6432 | 23,2 | 2 | NA | Neutral |
| TTN | rs2288569 | 2:179458591:C:T | NSM | 0,198 | 0,145342 | 0,052658 | 0,854648 | 24,7 | 2 | 0,479494362 | Deleterious |
| TTN | rs2288569 | 2:179458591:C:T | NSM | 0,198 | 0,145342 | 0,052658 | 0,854648 | 24,7 | 2 | NA | Deleterious |
| TTN | rs2303838 | 2:179444939:C:T | NSM | 0,223 | 0,174237 | 0,048763 | 0,825763 | 16,89 | 2 | 0,511611545 | Neutral |
| TTN | rs2303838 | 2:179444939:C:T | NSM | 0,223 | 0,174237 | 0,048763 | 0,825763 | 16,89 | 2 | NA | Neutral |
| TTN | rs2627037 | 2:179606538:G:A | NSM | 0,074 | 0,074158 | -0,000158 | 0,925842 | 21,9 | 2 | 0,375507967 | Neutral |
| TTN | rs2627037 | 2:179606538:G:A | NSM | 0,074 | 0,074158 | -0,000158 | 0,925842 | 21,9 | 2 | 0,376922289 | Neutral |
| TTN | rs2627043 | 2:179582537:G:T | NSM | 0,233 | 0,203063 | 0,029937 | 0,796937 | 22,7 | 2 | 0,37003315 | Neutral |
| TTN | rs35813871 | 2:179650408:G:A | NSM | 0,233 | 0,237909 | -0,004909 | 0,762091 | 22,4 | 2 | 0,488451209 | Deleterious |
| TTN | rs36051007 | 2:179545859:C:T | NSM | 0,243 | 0,302621 | -0,059621 | 0,697379 | 13,05 | 2 | NA | Neutral |
| TTN | rs3731746 | 2:179430997:G:A | NSM | 0,223 | 0,165749 | 0,057251 | 0,834221 | 22,5 | 2 | 0,501977067 | Deleterious |
| TTN | rs3731746 | 2:179430997:G:A | NSM | 0,223 | 0,165749 | 0,057251 | 0,834221 | 22,5 | 2 | 0,595561276 | Deleterious |
| TTN | rs3731746 | 2:179430997:G:A | NSM | 0,223 | 0,165749 | 0,057251 | 0,834221 | 22,5 | 2 | NA | Deleterious |
| TTN | rs3731749 | 2:179406191:C:T | NSM | 0,193 | 0,144008 | 0,048992 | 0,855992 | 25,5 | 2 | 0,472178479 | Deleterious |
| TTN | rs3731749 | 2:179406191:C:T | NSM | 0,193 | 0,144008 | 0,048992 | 0,855992 | 25,5 | 2 | NA | Deleterious |
| TTN | rs3829746 | 2:179427536:T:C | NSM | 0,267 | 0,226525 | 0,040475 | 0,773475 | 22 | 2 | 0,382623153 | Neutral |
| TTN | rs3829746 | 2:179427536:T:C | NSM | 0,267 | 0,226525 | 0,040475 | 0,773475 | 22 | 2 | 0,388016078 | Neutral |
| TTN | rs3829746 | 2:179427536:T:C | NSM | 0,267 | 0,226525 | 0,040475 | 0,773475 | 22 | 2 | NA | Neutral |
| TTN | rs3829747 | 2:179397561:C:T | NSM | 0,193 | 0,14416 | 0,04884 | 0,85584 | 25,2 | 2 | 0,459772507 | Deleterious |
| TTN | rs3829747 | 2:179397561:C:T | NSM | 0,193 | 0,14416 | 0,04884 | 0,85584 | 25,2 | 2 | 0,477549911 | Deleterious |
| TTN | rs3829747 | 2:179397561:C:T | NSM | 0,193 | 0,14416 | 0,04884 | 0,85584 | 25,2 | 2 | NA | Deleterious |
| TTN | rs6723526 | 2:179498042:T:C | NSM | 0,079 | 0,107524 | -0,028524 | 0,892476 | 23,6 | 2 | 0,490594059 | Deleterious |
| TTN | rs6723526 | 2:179498042:T:C | NSM | 0,079 | 0,107524 | -0,028524 | 0,892476 | 23,6 | 2 | 0,500231471 | Deleterious |
| TTN | rs6723526 | 2:179498042:T:C | NSM | 0,079 | 0,107524 | -0,028524 | 0,892476 | 23,6 | 2 | NA | Deleterious |
| TTN | rs72648998 | 2:179575511:C:T | NSM | 0,05 | 0,055601 | -0,005601 | 0,944399 | 23,1 | 2 | NA | Neutral |
| TTN | rs744426 | 2:179436020:G:A | NSM | 0,198 | 0,143718 | 0,054282 | 0,856282 | 23,8 | 2 | 0,457484084 | Deleterious |
| TTN | rs744426 | 2:179436020:G:A | NSM | 0,198 | 0,143718 | 0,054282 | 0,856282 | 23,8 | 2 | 0,522537827 | Deleterious |
| TTN | rs744426 | 2:179436020:G:A | NSM | 0,198 | 0,143718 | 0,054282 | 0,856282 | 23,8 | 2 | NA | Deleterious |
| TTN | rs922984 | 2:179615887:T:C | NSM | 0,931 | 0,928809 | -0,002191 | 0,071191 | 11,74 | 2 | 0,432056651 | Neutral |
| TTN | rs9808377 | 2:179421694:A:G | NSM | 0,267 | 0,224486 | 0,042514 | 0,775514 | 12,6 | 2 | 0,254387605 | Neutral |
| TTN | rs9808377 | 2:179421694:A:G | NSM | 0,267 | 0,224486 | 0,042514 | 0,775514 | 12,6 | 2 | 0,281592154 | Neutral |
| TTN | rs9808377 | 2:179421694:A:G | NSM | 0,267 | 0,224486 | 0,042514 | 0,775514 | 12,6 | 2 | NA | Neutral |
| XRN1 | rs1802904 | 3:142168331:C:T | UTR | 0,851 | 0,855129 | 0,004129 | 0,144871 | 17,22 | 1 | NA | NA |
| XRN1 | rs35351308 | 3:142141496:G:A | NSM | 0,025 | 0,016331 | 0,008669 | 0,983669 | 23,3 | 1 | 0,451343449 | Deleterious |
| XRN1 | rs73238159 | 3:142078759:C:T | NSM | 0,114 | 0,129277 | -0,015277 | 0,870723 | 25,4 | 1 | 0,409708481 | Deleterious |
| ZCCHC2 | rs866334998 | 18:60242678:G:T | NSM | 0,009901 | 0,000039 | 0,009862 | 0,999961 | 25,9 | 2 | 0,566640655 | NA |

**Supplementary Table S2.** Comparison of allele frequency of (rs7737765; H148Y) between IRIS1 cohort and gnomAD

|  | IRIS 1 | gnomAD (control) |
| --- | --- | --- |
| REF allele count | 168 | 1045683 |
| ALT allele count | 34 | 133271 |
|  | 202 | 1 178 954 |

Minor allele frequency (MAF) IRIS 1 cohort: 0.168
MAF gnomAD: 0.1130

**RESULT of statistical testing:**

Odds-ratio OR_REF_: 1.58794

P-value Fisher exact test: 0.01889

**Supplementary table S3.**
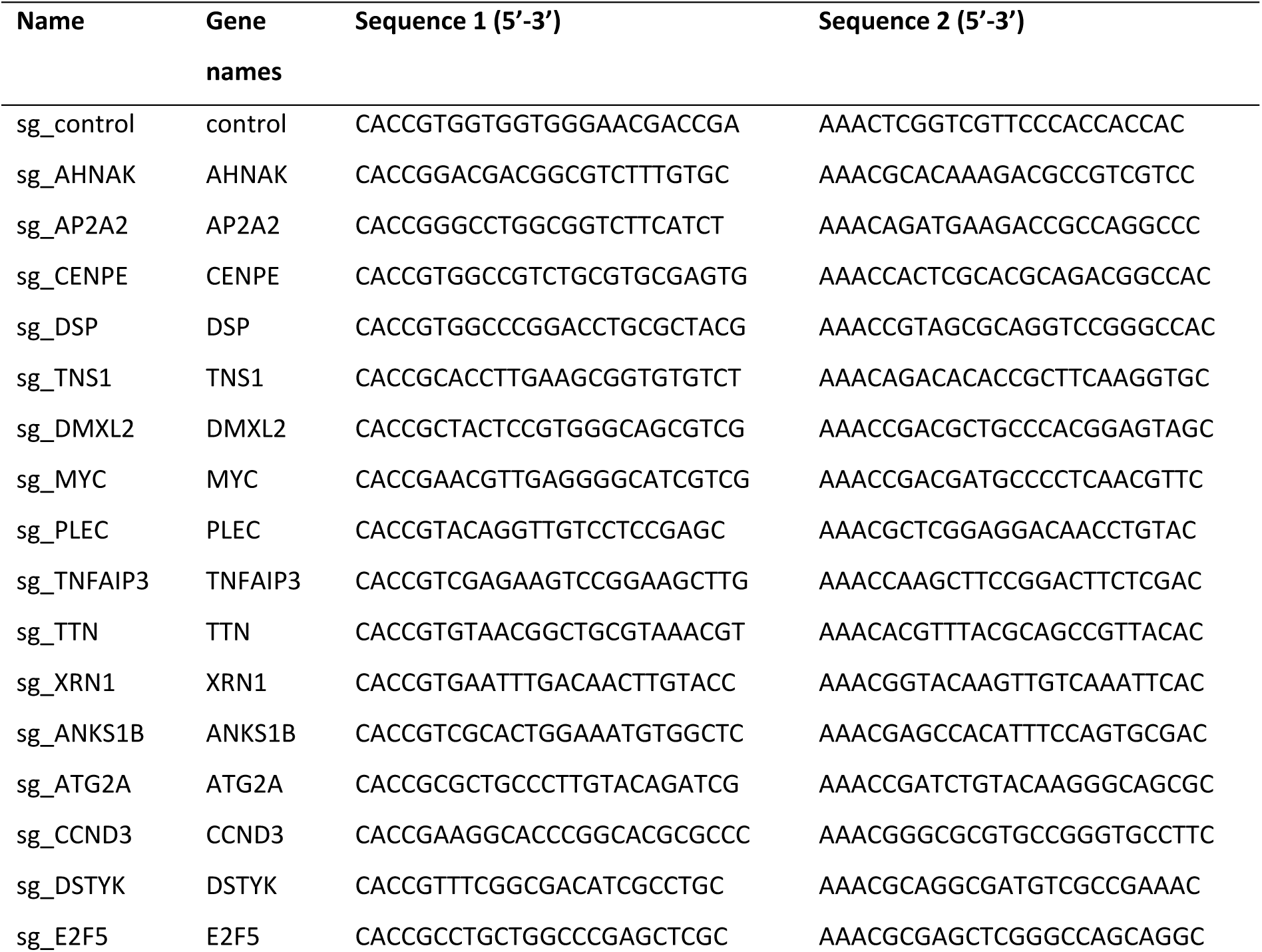
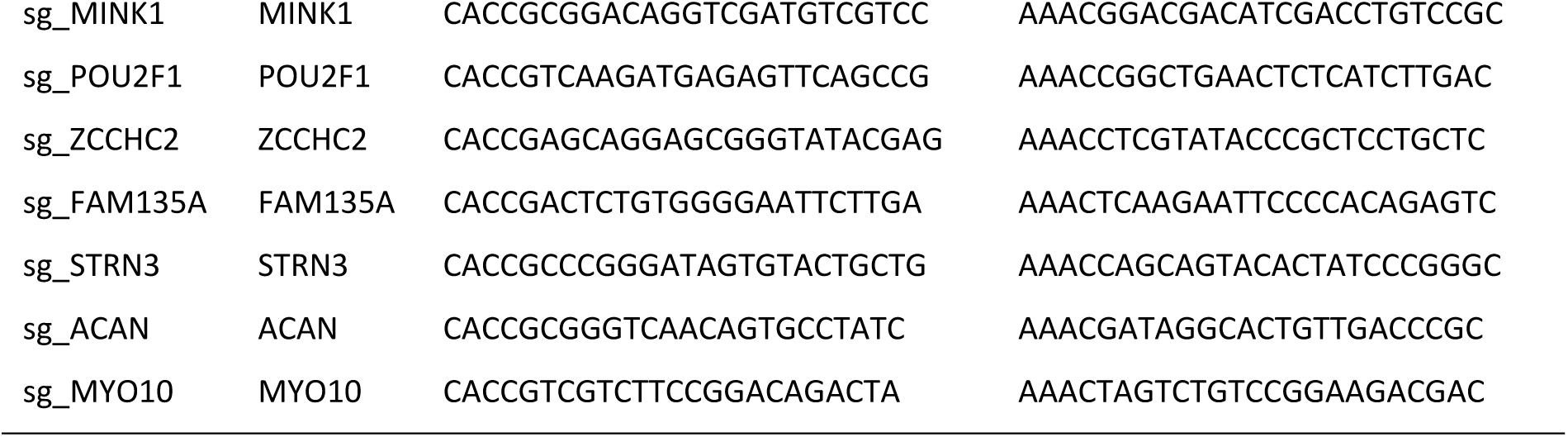
Oligonucleotides containing sgRNA sequences of candidate genes

**Supplementary table S4.**
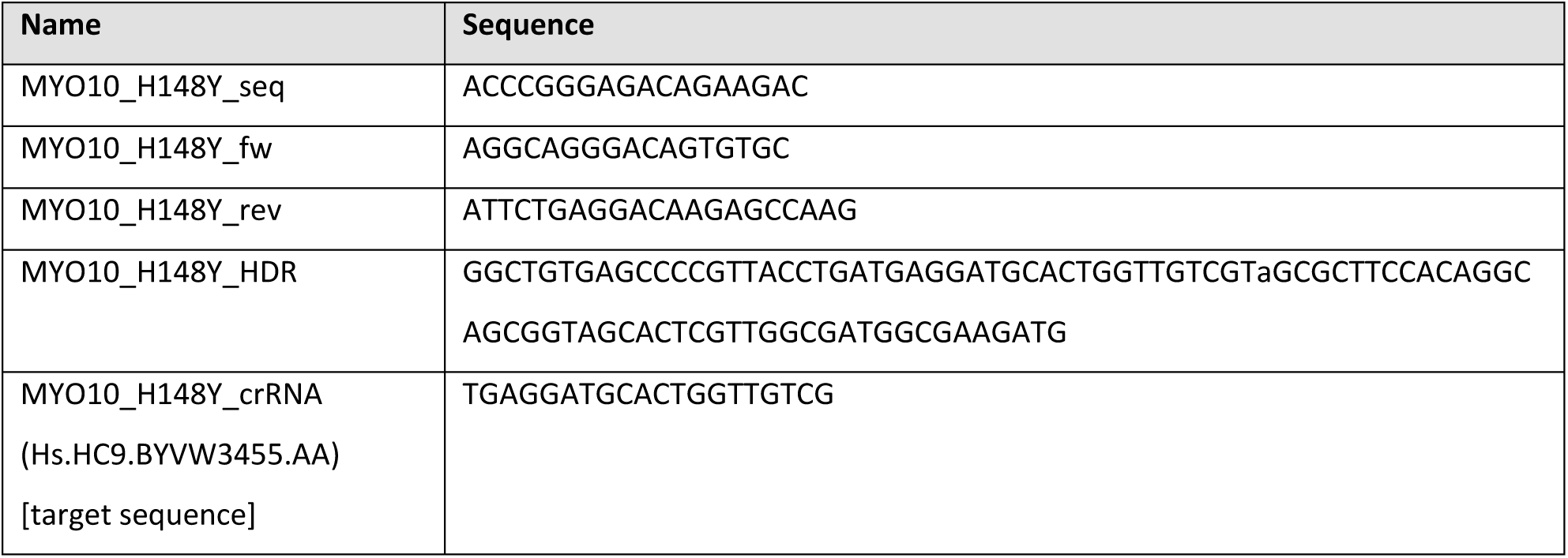
Oligonucleotides for CRISPR knock-in

